# Epigenetic reprogramming by 5-aza-dC alters gene methylation and suppresses aggression in founding queens of the California harvester ant *Pogonomyrmex californicus*

**DOI:** 10.1101/2025.09.11.675510

**Authors:** Tania Chavarria-Pizarro, Mohammed Errbii, Benne Köster, Mjtia Möhleke, Lukas Schrader, Juergen Gadau

## Abstract

Epigenetic mechanisms are increasingly recognized as key drivers of phenotypic plasticity in social insects, where the caste and function of an individual is usually not determined by its genotype. These mechanisms may also regulate variation in social behavior or interactions and ultimately social organization. We experimentally manipulated genome-wide DNA methylation in founding queens of *Pogonomyrmex californicus* using the DNA methylation inhibitor decitabine (5-aza-20-deoxycytidine), to investigate the effect of DNA demethylation on aggressive interactions during colony founding in *P. californicus*, leading either to colonies with single (i.e., monogyny) or multiple queens (i.e., polygyny). Genome-wide methylation profiling showed widespread gene hypomethylation in treated queens, including those encoding the DNA methyltransferases DNMT1 and DNMT3. Furthermore, treated queens exhibited significantly reduced aggression in queen-queen interactions compared to untreated ones. Interestingly, genes linked to energy metabolism showed increased methylation in aggressive founding queens compared to non-aggressive queens. Our results highlight a potential epigenetic basis for behavioral and social variation, contributing to the understanding of social niche polymorphism in ants.

## Introduction

An individual’s phenotype is the result of the interaction between its genotype and environment (Wong et al. 2005; Pozo et al. 2021). A key feature of this interaction is **phenotypic plasticity**—the ability of a single genotype to produce different phenotypes in response to environmental conditions. At its core, phenotypic plasticity is mediated by changes in gene expression (Holliday 1975; Jablonka & Lamb 2002; Richards 2006; Bird 2007; Dean & Maggert 2015), which are often regulated by epigenetic mechanisms such as DNA methylation, histone modification, chromatin remodeling, and non-coding RNAs (Grant-Downton & Dickinson 2005; Berger 2007). Epigenetic modifications are molecular changes that allow organisms to respond rapidly, reversibly, and sometimes long-term to environmental changes (Wong et al. 2005; Pozo et al. 2021). In animals, DNA methylation commonly occurs at CpG sites and is controlled by two main enzymes: DNA methyltransferase 3 (DNMT3), which establishes new methylation patterns, and DNMT1, which maintains existing ones (Goll & Bestor 2005; Cheng & Blumenthal 2008; Kim et al. 2008). In insects, methylation is typically concentrated in gene bodies (Feng et al. 2010; Lyko et al. 2010; Bonasio et al. 2012).

In eusocial Hymenoptera (ants, bees, and wasps), DNA methylation has been implicated in both behavioral plasticity and caste differentiation, with most species retaining functional DNMT1 and DNMT3 genes (Yan et al. 2014; Yan et al. 2015; Okwaro & Korb 2023). These insects exemplify extreme phenotypic plasticity, with highly divergent reproductive, morphological, and behavioral castes arising from a shared genome (Kucharski et al. 2008; Herb et al. 2012; Yan et al. 2015). Still, the precise role of DNA methylation in regulating social behavior remains unresolved. While some studies report associations between methylation patterns and caste or behavioral traits (Elango et al. 2009; Bonasio et al. 2012; Patalano et al. 2015), others find minimal or inconsistent effects (Libbrecht et al. 2016; Standage et al. 2016).

Beyond caste differentiation, plasticity in eusocial systems also manifests in colony structure. Social organization in ants, for example, can vary dramatically—from monogynous colonies with a single queen to polygynous ones with multiple reproductive queens (Keller 1993; Boomsma et al. 2014; Saltz et al. 2016). These alternative strategies may represent adaptive “social niches,” analogous to ecological niches, in which individuals maximize their inclusive fitness through different social roles (Saltz et al. 2016; Kaiser et al. 2024). Variation in queen number has been observed within and between species, including *Messor pergandei*, *Acromyrmex versicolor*, and the *Formica rufa–F. polyctena* species complex (Cahan & Julian 1999; Gyllenstrand, Seppä & Pamilo 2004; Helms & Cahan 2012).

The California harvester ant, *Pogonomyrmex californicus*, provides a particularly informative case. Its populations exhibit both monogynous and polygynous colony structures (Johnson 2004; Overson et al. 2014). Colonies can be initiated by a single queen (haplometrosis) or cooperatively by multiple queens (pleometrosis), resulting in enduring differences in social organization (Keller 1993; Boomsma et al. 2014). These strategies are accompanied by distinct behavioral patterns: haplometrotic queens are aggressive toward rival queens, while pleometrotic queens are more tolerant (Clark & Fewell 2014; Overson et al. 2014).

Remarkably, neighboring populations of *P. californicus* often show stark differences in founding behavior—some are nearly entirely monogynous, while others are predominantly polygynous (Johnson et al. 2004; Overson et al. 2014). Laboratory studies demonstrate that these behavioral strategies are both stable and heritable (Rissing et al. 2000; Clark & Fewell 2014). Transcriptomic data further support this distinction: aggressive queens show elevated expression of genes involved in metabolism, immunity, and neural function, whereas tolerant queens upregulate chemosensory genes (Helmkampf et al. 2016).

This behavioral and genetic divergence positions *P. californicus* as a powerful model for exploring the epigenetic underpinnings of social plasticity. Genomic analyses have identified an 8 Mb supergene-like region associated with social structure that includes DNMT3, while DNMT1 shows signatures of selection in pleometrotic populations (Erbii et al. 2024). Methylome profiling reveals low but functionally relevant genome-wide methylation (∼3%), concentrated in gene bodies and positively correlated with expression (Chavarria-Pizarro et al. 2025). Methylation levels also vary across castes and developmental stages, with metabolically active individuals such as queens and pupae showing higher methylation levels— suggesting a plastic regulatory role in caste differentiation and behavior.

Cooperative nest founding in queens of *P. californicus* thus offers a valuable system for probing how epigenetic mechanisms, particularly DNA methylation, shape social interactions and niche construction. In this study, we used 5-aza-2′-deoxycytidine (5-aza-dC), a chemical inhibitor of DNA methylation, to experimentally manipulate methylation levels in founding queens. In Hymenoptera, 5-aza-dC—typically administered through sugar water—has been shown to produce striking phenotypic changes. Once taken up, 5-aza-dC is converted into 5-aza-dCTP and incorporated into replicating DNA, where it irreversibly binds to DNMT1, blocking maintenance methylation and causing passive demethylation through cell divisions (Christman 2002; Hagemann et al. 2011; Ramos et al. 2015; Seelan et al. 2018). Previous experiments illustrate the behavioral impact of this compound: in *Nasonia vitripennis*, it biased sex ratios toward males (Cook et al. 2015), while in *Bombus terrestris*, it accelerated colony development and increased cooperation without disrupting social hierarchies (Pozo et al. 2021).

This study aimed to determine how experimental disruption of DNA methylation affects queen behavior in *Pogonomyrmex californicus*, with a focus on aggression during nest founding. Specifically, we investigated whether treatment with 5-aza-2′- deoxycytidine (5-aza-dC) alters aggressive behavior and whether distinct DNA methylation patterns are associated with behavioral phenotypes (aggressive vs. non-aggressive) and chemical treatment (treated vs. non-treated).

Our results provide experimental evidence that DNA methylation plays a role in regulating queen behavior. Queens treated with 5-aza-dC exhibited significantly reduced aggression compared to untreated controls. Methylation profiling revealed global hypomethylation in treated individuals, including in key regulatory genes such as DNMT1 and DNMT3. Furthermore, the most aggressive individuals (aggressive control queens) showed elevated methylation in genes involved in energy metabolism—aligning with known links between metabolic gene expression and aggression. These findings support a potential epigenetic basis for behavioral plasticity and social variation in this species’ social niche polymorphism.

## Methods

### Experimental setup 2023

In 2023 120 founding queens were collected directly after their nuptial flight at Salt River, Phoenix Arizona (33.55034 N, − 111.64453 W) between May and June. Queens were put individually in separate petri dishes and kept at 26 °C on a diet of sugar water and seed.

A stock solution of 5-aza-20-deoxycytidine (decitabine) was made by dissolving 5 mg of decitabine (Sigma Aldrich) in 2 ml of distilled water solution (10mM). Decitabine was added to sugar water (0.0925% v/v) (Amarasinghe et al. 2014; Pozo et al. 2021) and fed to 60 queens for 10 days ad libitum. The control group (60 queens) was fed with standard sugar water for 10 days, also at 0.0925% v/v. Solutions were provided to each queen daily throughout 10 days by putting five drops of the solution on a cotton ball. Pre-treatment with no sugar-water for three days ensured that both treatment and control colonies readily were hungry enough to drink the sugar-water during the feeding period.

After the feeding period (10 days) founding queens were placed in pairs in standardized observation nests containing sterilized soil from the collection site and grass seeds as food. Only specimens of similar body weight (within 10% of each other’s weight, mean = 13.5 mg, total range = 9.5– 20 mg) were paired to avoid effects of body size on social interaction. Two experimental groups were established in 30 replicates each: control pairs and treatment pairs. Overt aggressive behaviour, defined as grappling, biting and chasing, was recorded for 3 days by semi-continuous scan sampling. Each nest was observed for approximately 30 s before moving on to the next one (see Holbrook et al. 2009). On average, 36 rounds of observations were completed per pair for a total of 108 observations. We also observed non-aggressive behaviors, like receiving aggression, antennation, resting, digging and activity.

After the end of the three-day observation period, all living queens were collected and conserved in ethanol.

### Experimental set up 2024

We repeated the behavioral experiment in the summer of 2024 to assess whether the 10-day isolation period—during which queens were fed the solution of 5-aza-20-deoxycytidine (decitabine) (treatment) or the sugar-water (control)—had any effect on the behavioral patterns observed in 2023. In particular, whether this waiting period before colony founding significantly reduced aggressive interactions. 90 founding queens were collected directly after their nuptial flight from the same site and population (Salt River, Phoenix Arizona 2023 (33.55034 N, − 111.64453 W). Queens were housed individually in petri dishes at 26□°C and provided with water only for three days. After this acclimation period, 46 specimens were paired based on body weight (within 10%, mean = 13.5 mg, range = 9.5–20 mg) to control for size-related behavioral effects. For the behavioral analyses they were treated and analysed exactly like in 2023. Behavioral observations began immediately and were recorded over six days. These constituted the immediate control group. Between 13 and 17 observation rounds were completed daily, for a total of 112 observations. The second group of queens were paired after a 10-day feeding period. Pairing was done exactly like in the immediate group This setup mirrored the untreated group in our 2023 experiment. Behavioral observations—both aggressive and non-aggressive—were conducted over four days, with 17 rounds per day, totaling 68 observation rounds (Fig 1).

**Figure 1.**
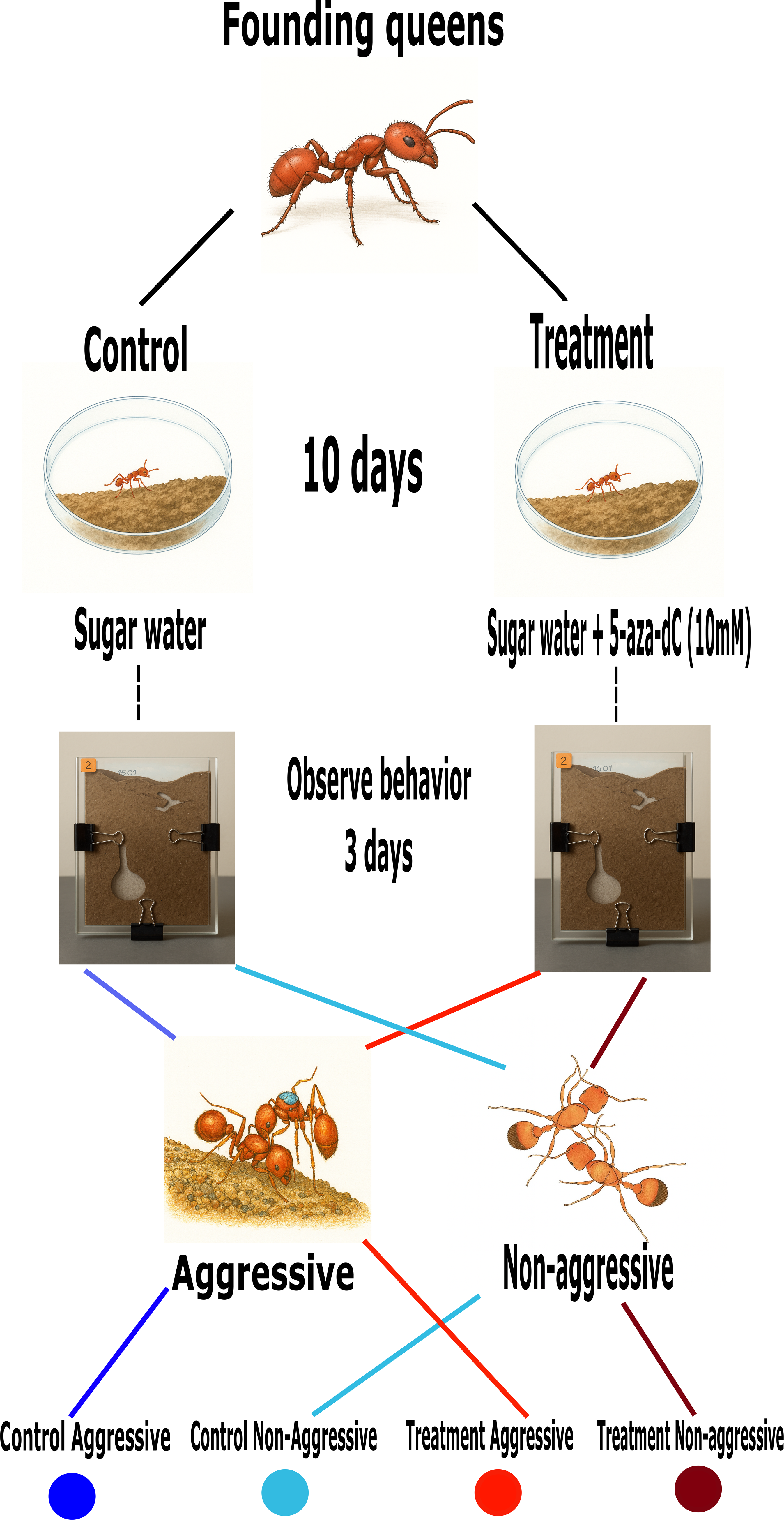
Experimental design of the epigenetic reprograming. 120 founding queens were collected and put individually in separate petri dishes and kept at 26 °C on a diet of sugar water and seed. A stock solution of 5-aza-20-deoxycytidine (decitabine) was added to sugar water (0.0925% v/v) and fed to 60 queens for 10 days ad libitum. The control group (60 queens) was fed with standard sugar water for 10 days, also at 0.0925% v/v. After the feeding period (10 days) founding queens were placed in pairs in standardized observation nests. Two experimental groups were established in 30 replicates each: control pairs and treatment pairs. Overt aggressive behaviour was recorded for 3 days by semi-continuous scan sampling. An individual was considered aggressive if it displayed more than 10 aggressive behaviors during the whole observation period. Based on this criterion, we classified individuals into four categories: Control aggressive (CA) (blue), Control non-aggressive (CNA) (light-blue), Treatment aggressive (TA)(red), Treatment non-aggressive (TNA)(dark-red).

After the end of the three-day observation period, queen specimens were collected and stored in 100 % ethanol. These generated four cohorts across two years, whereby two groups, untreated and pairing delayed for 10 days, overlap (Fig 1).

### Behavioural analysis

We focused our observations on six behavioral categories: 1. Aggressive behavior (biting, grabbing, chasing) 2. Receiving aggressive behavior 3. Activity (movement) 4. Resting 5. antennation and 6. Digging.

We recorded the occurrence of each behavior during each observation round and then calculated the average number of occurrences per observation hour. To compare behavioral averages across treatments and sampling years, we used a Kruskal-Wallis test due to the non-normal distribution of the data. When significant differences were found, we performed Dunn’s post hoc test for pairwise comparisons.

In 2023, we analyzed two groups: treatment and control. In 2024, the groups were immediate and delayed (same as control 2023). For the 2023 dataset, we further categorized individuals based on their level of aggression. An individual was considered aggressive if it displayed more than 10 aggressive behaviors during the whole observation period. Based on this criterion, we classified individuals into four categories:

1. Control aggressive (CA)
2. Control non-aggressive (CNA)
3. Treatment aggressive (TA)
4. Treatment non-aggressive (TNA)

Finally, we made a ranking of aggression of each individual in 2023. For this we took the total number of aggressive behavior of each individual and divided it for the maximum number of aggressive behavior found in an individual (Fig 1).

### DNA isolation and sequencing

DNA was extracted from whole bodies of frozen or ethanol-preserved individuals of *P. californicus* with the Qiagen DNeasy Blood and Tissue kit. A Qubit BR assay was used to assess DNA quantity, followed by 1 % agarose gel electrophoresis to confirm the presence of high molecular weight DNA (>10 kb). We prepared DNA libraries of 20 queens (10 treated and 10 nontreated) for sequencing using the Oxford Nanopore Technologies (ONT) MinION platform according to the user manual as also described in Chavarria-Pizarro et al. 2024. Basecalling and demultiplexing was performed with ont-guppy/6.4.8-CUDA-11.7.0 using the appropriate base calling model (dna_r9.4.1_450bps_hac.cfg) and default parameters (More details in https://github.com/TaniaChP79/P.cal-methylome). Variant calling was performed with **Longshot v0.4.3** (Edge & Bansal, 2019). Individuals with poor sequencing quality were excluded by calculating the number of SNPs per sample and removing those with fewer than 10,000 SNPs. The resulting VCF files were merged using **VCFtools v0.1.16** (Danecek et al., 2011) with the -F and --output-ref options to generate a list of candidate variant sites, ensuring regenotyping was restricted to predefined SNP positions. Variants with a sequencing depth greater than five were removed using **bcftools**. Principal component analysis (PCA) was then performed to assess genetic differences among queens from different treatments (More details in https://github.com/TaniaChP79/P.cal-methylome). Finally, we used 16 queens from the field season of 2024 (10 non-delay and 6 delay) to generate WGBS libraries to compare between treatments.

### Genome Assemblies and Annotations

We used the latest genome assembly and annotation for *P. californicus* (Pcal.3.1) (Errbii et al. unpublished) based on combined ONT long-read sequencing. This annotation included 15,899 protein-coding genes and had a TE proportion of 22.79 % (Errbii et al. unpublished).

### CpG Methylation Calling

We used Nanopolish v0.13.3 (Simpson et al. 2017) with default parameters to detect CpG methylation in ONT data (Liu et al. 2021; Yuen et al. 2021; Chavarria-Pizarro et al. 2024). Nanopolish indexes were generated from nanopore reads and aligned to the *P. californicus* reference genome (Pcal 3.1) using minimap2 v2.14 (Li 2018). Details are provided in Chavarria-Pizarro et al. (2025).

Nanopolish distinguishes 5-methylcytosine from unmethylated cytosines by detecting signal disruptions in raw ONT FAST5 data and calculates log-likelihood ratios for base modifications. Positive values indicate evidence of methylation.

To quantify methylation, we used the helper script calculate_methylation_frequency.py (Simpson et al. 2017), which outputs methylation frequency at genomic coordinates. We filtered methylation calls to retain only CpG motifs with a minimum read coverage of 10. Regions with less than 10% of CpG sites showing methylation were excluded (Perez et al. 2023).

We then created a BED file with all genomic CpG sites and mapped them to the corresponding methylation coverage and frequency using the bedtools map function (Quinlan & Hall 2010). This mapping was performed for both transcriptome and transposable element (TE) annotations (Errbii et al., unpublished).

### Inferences of gene body methylation

To exclude low-coverage data, we applied the commonly used threshold of at least 10 methylation calls per k-mer per sample (Perez et al. 2023).

For the analysis of average DNA methylation across annotated genes, we examined CpG methylation within 4 kb windows both upstream of the transcription start site (TSS) and downstream of the transcription termination site (TTS). We then calculated average methylation levels across these regions and within gene bodies by iterating over all annotated genes. We used a Kruskal–Wallis test to determined if there were significant differences in average methylation levels among the groups: (aggressive-control (CA), aggressive-treatment (CA), nonaggressive-control (CNA), nonaggressive-treatment (TNA) (See above Behavioral analysis for more details).

To determine whether coding regions exhibited higher methylation than non-coding regions, we performed a binomial test (Takuno & Gaut 2012) on CpG sites, comparing observed methylation to background levels. Because methylation could vary by treatment and behavior effect, tests were conducted separately for each group (See above Behavioral analysis for more details) (Bewick et al. 2016; Muyle et al. 2021). P-values were corrected for multiple testing using the Benjamini-Hochberg method (1995) within each treatment.

Genes were then classified into two categories based on adjusted p-values:

1. **Gene Body Methylated (GbM):** CpG methylation significantly higher than expected.
2. **Unmethylated (UM):** CpG methylation not significantly different from background.

Finally, we performed Gene Ontology (GO) term enrichment analysis on all significantly gene-body methylated genes using the **topGO** R package (v2.56; Alexa & Rahnenfuhrer 2024) in R v4.5.0. Tests for overrepresented GO terms in **Cellular Component, Biological Process and Molecular function** were calculated using **Fisher’s exact test** with hierarchical correction (weight01) which generates a table of the **top 200** most enriched GO terms sorted by significance p < 0.05.

### Differential Gene body methylated frequencies

To determine whether aggressive behavior and treatment influence DNA methylation frequencies in specific genes, we conducted pairwise comparisons using the Wilcoxon rank-sum test between Control and Treatment groups. We performed six comparisons:

1. Control Aggressive (CA) vs. Control Non-Aggressive (CNA)
2. CA vs. Treatment Aggressive (TA)
3. CA vs. Treatment Non-Aggressive (TNA)
4. CNA vs. TA
5. CNA vs. TNA
6. TA vs. TNA

We hypothesized that comparisons involving aggressive individuals would reveal a greater number of shared body methylated genes (GBM) than comparisons involving non-aggressive individuals. Given that the treatment reduced aggressive behavior, we expected fewer shared GBM genes in comparisons between aggressive and non-aggressive categories (e.g., CA vs. CNA, CA vs. TA, CNA vs. TA, TA vs. TNA), and more shared GBM genes between non-aggressive groups (e.g., CNA vs. TNA). The Benjamini-Hochberg (1995) correction was applied to the p-values (p.adjust_value, method = “BH”) to control the false discovery rate (FDR). This is important when performing multiple comparisons to reduce the likelihood of false positives. The log2 fold change was calculated as the difference between the log-transformed means of the Treatment and Control conditions. We considered that a gene was significantly differentially methylated between treatments if a gene had an adjusted p-value (adj_p_value) less than 0.05, with an absolute log2 fold change greater than 0.3, indicating that there was meaningful change in methylation between conditions of at least 30%. We used enhanced volcano plots to illustrate the test comparisons.

Finally, we examined the link between each individual’s aggression rank (see Behavioral Analysis for details) and the methylation frequencies of genes that showed significant differences in methylation across comparisons using a spearman correlation test, to determine whether these frequencies increased or decreased with aggression levels. Spearman correlation P-values were corrected for multiple testing using the Benjamini-Hochberg method (1995).

### DNMT1 and DNMT3 methylation frequency

We calculated methylation frequency along treatments of the maintenance (DNMT1) and de novo (DNMT3) DNA methyltransferases using the data generated with ONT sequencing.

### Whole Genome Bisulfite sequencing

To assess whether the 10-day isolation period—during which queens were fed the solution of 5-aza-20-deoxycytidine (decitabine) (treatment) or the sugar-water (control)—had any effect on the DNA methylation changes that we observed in 2023. We compared DNA methylation frequencies of 16 queens from the field season of 2024 (10 non-delay and 6 delay) using WGBS data. The WGBS sequencing libraries were prepared by Novogene (Munich, Germany). In short, paired-end 150 bp bisulfite libraries were sequenced on the Illumina NovaSeq 6000 platform to a total of 35 million paired-end reads (nuclear coverage >20X). Reads were trimmed with trimmomatic v0.30 (Bolger, Lohse & Usadel 2014) (parameters: leading = 10, trailing = 10, minlen = 50) and processed with bismark v0.22.3 (Krueger & Andrews 2011) to compute per-base-pair methylation frequencies. For WGBS data, we calculated methylation sites of 16 queens from the field season of 2024 (10 non-delay and 6 delay) using the R package *methylKit v1.15.3* and R version 4.5.1 (Akalin et al. 2012). The percentage of methylated cytosines was calculated at a given site from the methylation ratios created by the software BSSeeker2 and complementary CpG dinucleotides were merged. For the analysis of average DNA methylation across annotated genes as we did with the ONT data (See above for more details). We then calculated average methylation levels across these regions and within gene bodies by iterating over all annotated genes. We used a Wilcoxon rank-sum test to determine if there were significant differences in average methylation levels along the gene region among the groups delay and non-delay groups.

### Differential Gene body methylated frequencies with WGBS for data 2024

To determine whether the 10-day isolation period had any effect on the DNA gene methylation changes that we observed in 2023 (see above for more details), we conducted pairwise comparisons using the Wilcoxon rank-sum test between delay and non-delay groups. We hypothesized that comparisons involving delay and non-delay groups will be mininum or not statistically difference. We applied the Benjamini-Hochberg (1995) correction to the p-values and calculate the log2 fold change was calculated as the difference between treatments (see above for more details). Then we used enhanced volcano plots to illustrate the test comparisons.

## Results

### Behavioral assays

During the observation period in 2023 and out of the 90 founding associations observed, including 45 control and 45 treated, a total of 49 pairs showed at least one instance of aggressive behavior, which was usually fighting, during the 3-day observation period. Control queens displayed aggressive behaviour significantly more frequently than treated queens, both in terms of the number of nests in which aggressive behaviour was observed (Fig. 2, Kruskall-Wallis test, p = 0.01), and in the absolute number of aggressive acts (Fig. 2, Appendix 1). In addition, treated queens rest more (Fig. 2, Kruskall-wallis test, p = 0.005) than control queens and have the tendency to be less active (movement) (Fig. 2, Kruskall-Wallis test, p = 0.005). These results demonstrate that aggression decreased when queens were chemically treated with decitabine. Nonetheless, the number of nests in which aggressive interactions occurred was generally low in control and treated queen nests (Appendix 3-4), which was unexpected (see discussion).

**Figure 2.**
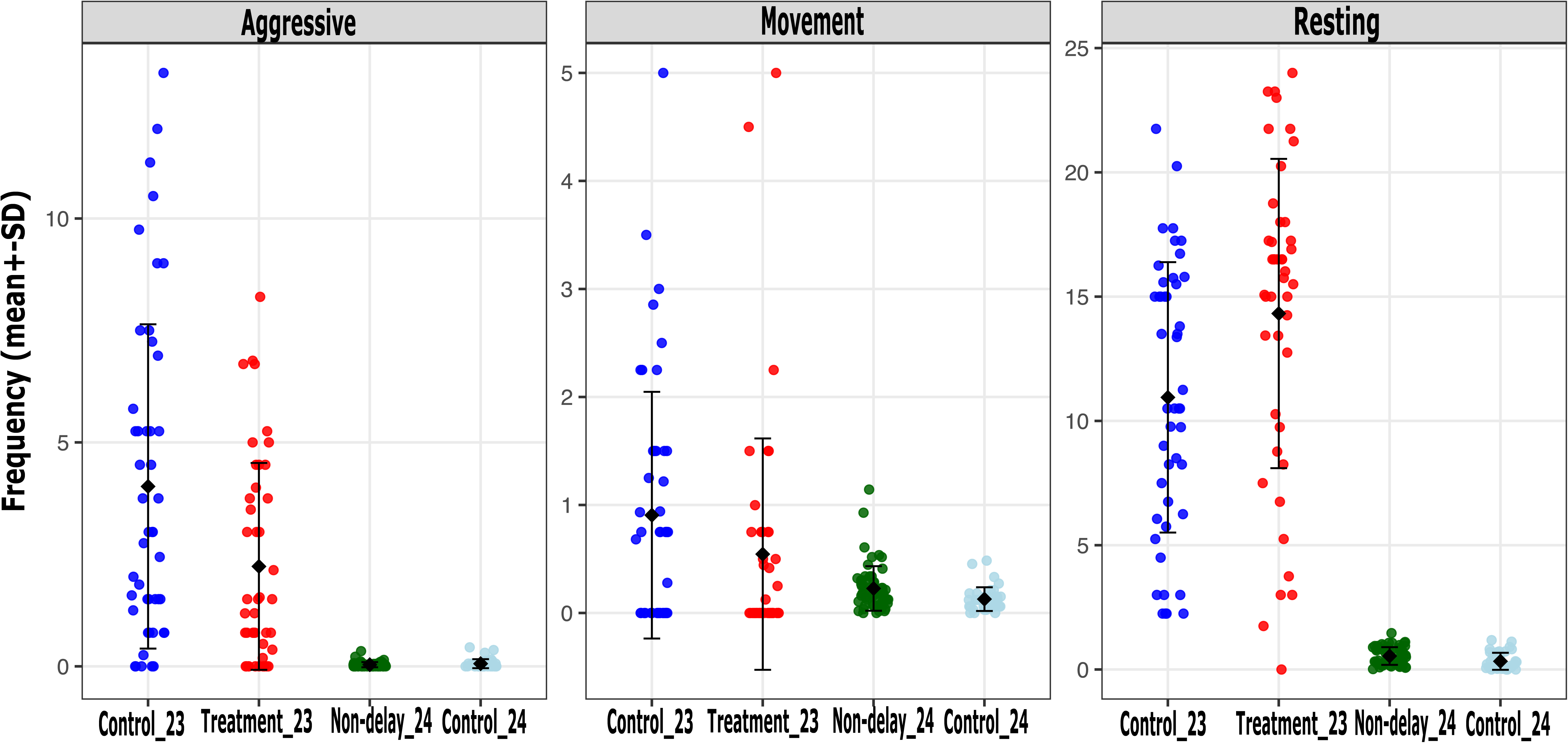
Average and standard error of the behavioral frequencies that show significant difference across treatments and sampling years. We used a Kruskal-Wallis test and Dunn’s post hoc test for pairwise comparisons due to the non-normal distribution of the data. In 2023, we had two groups: Isolation 23 (Control 23) and Treatment 23. In 2024, the groups were no non-delay 24 and delay (control 24).

As described in the Methods section, individuals were classified as aggressive if they displayed more than 10 aggressive behaviors, including grappling, biting, and chasing. However, aggression levels were notably reduced in aggressive individuals that received chemical treatment. For instance, the frequency of aggressive behaviors in Control Aggressive (CA) queens ranged from 10 to 108 events, whereas in Treatment Aggressive (TA) queens, the range was lower, from 10 to 66 events.

We repeated the behavioral experiment in the summer of 2024 to assess whether the 10-day isolation period—during which queens were fed the solution of 5-aza-20-deoxycytidine (decitabine) (treatment) or the sugar-water (control)—had any effect on the behavioral patterns observed in 2023, in particular made queens less aggressive. From the 60 founding associations including 30 control and 30 delayed (see methods for more details), a total of 53 pairs showed at least one instance of overt aggression (i.e. fighting) during the 3-day observation period. Control queens displayed similar aggressive behaviour then delayed queens (Fig. 2, Kruskal-Wallis test, P = 0.554). In addition, control queen rest more (Fig. 2, Kruskal-Wallis test, P = 0.0009) than delayed queens and delayed queens were less active (movement) (Fig. 2, Kruskal-Wallis test, P = 0.001)(Appendix 1-2).

### DNA methylation profiles with ONT between treatments

To detect DNA 5-methylcytosine in CpGs (5mCpG) in control and treated queens, we generated on average 18.3 million reads per adult sample with ONT sequencing (Table. 1). Genome-wide methylation levels varied among treatments with nonaggressive queens treated with decitabine showing the lowest percentage of methylated CpG sites (2.3 %), followed by aggressive treated queens (2.5%), control nonaggressive queens (2.6 %) and highest in control aggressive queens (2.8 %). In addition, the overall number of methylated and unmethylated genes differed between these samples (Table. 1). TE families also differed in methylation frequencies between the treatments, with high methylation levels in LINE elements in control queens compared with treated ones (Fig S1).

The DNA methylation profiles of *P. californicus* founding queens showed low levels of CpG methylation (2.5-3%) across treatments (Table 1), with the highest methylation levels observed in genic regions (9.5 %), followed by promoter regions (2.7 %), intergenic regions (2.6%?), and TEs being the least methylated (2.5 %) (Table 2). Average methylation for gene regions was highest for control aggressive and control nonaggressive queens and lowest for treated aggressive queens (Fig 2A). Methylation patterns within the genome differ between treatments particularly at the gene level, with the highest methylation levels found in the control aggressive queens (Table 2). A Kruskal–Wallis test revealed significant differences in average methylation levels among the four groups (χ² = 797.8, df = 3, p < 2.2 × 10⁻¹□).

**Table 1.** Nanopore ONT sequencing – queen methylation data per treatment and behavior (control:untreaded, treatment:treated). Average methylation was calculated as the average CG gene methylation across all genes. We calculated average DNA methylation for each treatment separaly, as methylation levels differ between these categories. A gene was considered methylated if had a significantly higher proportion of methylated cytosines than the genome-wide methylation average in each stage or caste. This was performed using a Binomial test and p-values were corrected for multiple tests.

| Treatment | N | Total called CpG sites | Methylated CpG sites | Average DNA methylation all CpG sites | Average DNA methylation frequency on genes (TSS-TTS) | no. methylated genes | no. Un-methylated genes |
| --- | --- | --- | --- | --- | --- | --- | --- |
| Control aggressive | 6 | 491,275,307 | 13,116,859 | 0.028 | 0.038 | 5300 | 9105 |
| Control nonaggressive | 6 | 338,142,274 | 8,209,837 | 0.026 | 0.038 | 5201 | 9204 |
| Treatment | 12 | 304,653,936 | 8,683,992 | 0.025 | 0.038 | 5095 | 9300 |

**Table 2.** Methylation frecuency patterns along the genome between treatment and behavior (average+ standard deviation) (control:untreaded, treatment:treated).

| Treatment | Exon | five_prime_<br>UTR | intron | intergenetic | mRNA | Promoter | TE | three_prime_<br>UTR | tRNA |
| --- | --- | --- | --- | --- | --- | --- | --- | --- | --- |
| Control<br>aggressive | 0.093±0.0<br>002 | 0.053±0.000<br>09 | 0.047<br>±0.000001 | 0.037<br>±0.000001 | 0.066<br>±0.0001 | 0.0421<br>±0.0008 | 0.042<br>±0.0008 | 0.066<br>±0.0004 | 0.056<br>±0.0022 |
| Control non<br>aggressive | 0.091±0.0<br>002 | 0.051±0.000<br>09 | 0.035<br>±0.000009 | 0.045<br>±0.000001 | 0.065<br>±0.0011 | 0.045<br>±0.0008 | 0.039<br>±0.0008 | 0.066<br>±0.0004 | 0.047<br>±0.0021 |
| Treated | 0.089±0.0<br>002 | 0.052±0.000<br>09 | 0.063<br>±0.00001 | 0.037±3.19<br>E+04 | 0.065<br>±0.0010 | 0.067<br>±0.0008 | 0.042<br>±0.0008 | 0.042<br>±0.0004 | 0.068<br>±0.0021 |

Subsequent pairwise Wilcoxon rank-sum tests (BH-adjusted p values) indicated that almost all group comparisons were significant. Specifically, both treatment groups (Aggressive and Non-Aggressive) showed significantly lower methylation compared to the corresponding control groups (p < 2 × 10⁻¹□). Moreover, the Control Aggressive and Control Non-Aggressive groups also differed significantly (p = 0.033). In contrast, no significant difference was detected between Treatment Aggressive and Treatment Non-Aggressive (p = 0.109). Principal components analyses of overall queens SNPs some level of structure (Fig S2; PC1: 15% and PC2: 13% variation respectively). Nonetheless, when we removed the outliers, we found that the dominant genome-wide variation captured by PC1/PC2 is **not** aggression status (Fig S3, PC1: 10% and PC2: 8% variation respectively).

Analyses of gene body-methylation revealed a high number of methylated genes in control aggressive queens, whereas a low number of methylated genes was detected in treated non-aggressive queens (Fig. 3A, Table 1). Approximately 77 % (4040) of the GBM genes were shared between all treatments (Fig. 3B). Enrichment analyses of these genes revealed functional categories associated with housekeeping processes, including DNA replication, translation and transcription factors, RNA processing, and genes corresponding to methyltransferases (Appendix 3).

**Figure 3.**
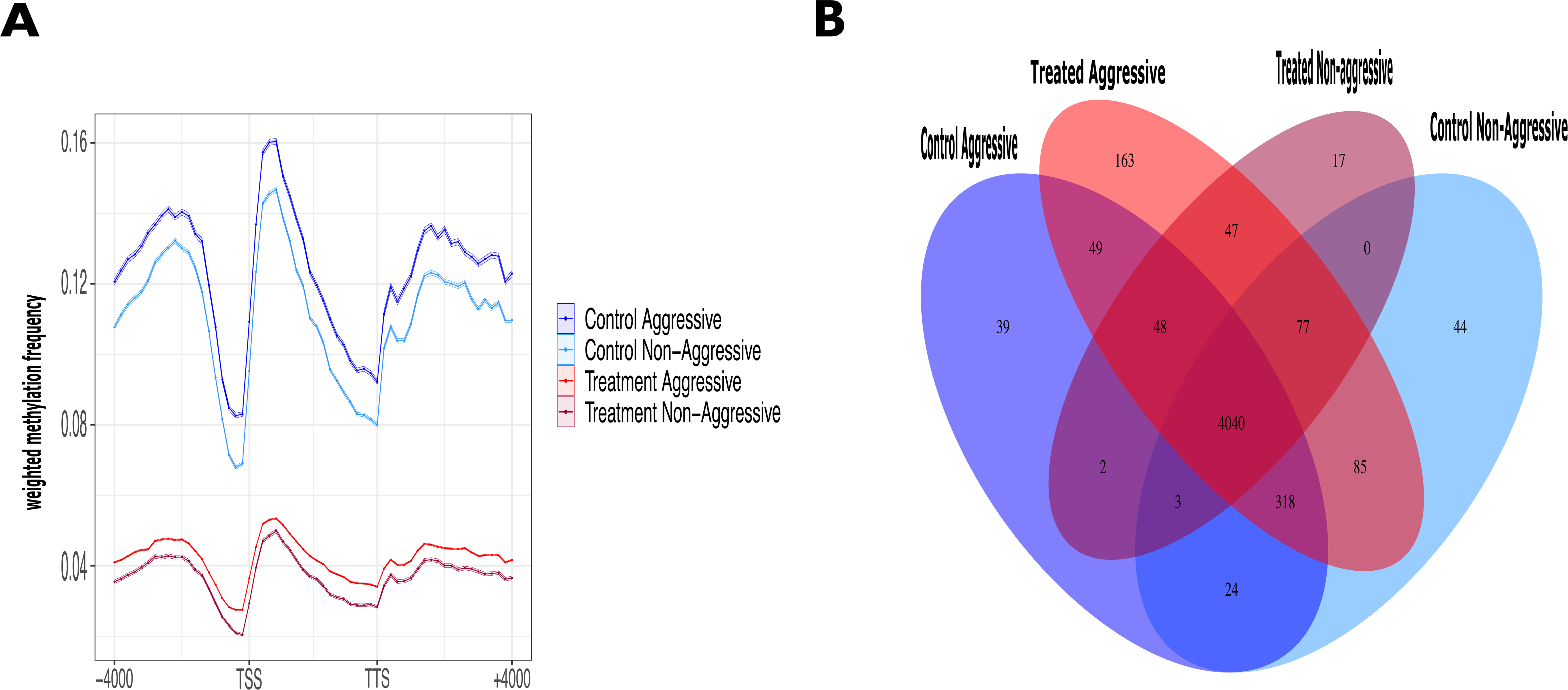
Gene methylation patterns in founding queens along behaviour differences (aggressive and non-aggressive) and treated individuals with decitabine (treatment) and nontreated (controls) obtain with the sequencing method ONT A) Weighted frequency methylation average among treatments (n=6 control aggressive, n=6 control non aggressive, n=12 treatment) looped over all genes region within a window of 4 kb upstream of the transcription start site (TSS) and a window of 4 kb downstream of the transcription end site (TTS) B) Venn diagram of Genes body methylated unique and share along treatments (control aggressive=dark blue, control non aggressive=lightblue, treatment=red).

We observed a low number of uniquely gene body methylated (GBM) genes in each treatment group (Fig. 3B). Contrary to our expectations, the highest number of shared (GBM) genes was found between Control Aggressive and Control Non-Aggressive individuals (345 genes), suggesting that chemical treatment has a stronger effect on GBM patterns than behavioral phenotype alone.

Nonetheless, aggression still appeared to influence GBM profiles. Control Non-Aggressive queens shared more GBM genes with treated individuals (77 genes) than Control Aggressive queens did (48 genes), indicating some behavioral influence on methylation patterns (Fig. 3B).

Genes uniquely methylated in Control Aggressive queens (CA) were primarily associated with transcription and intracellular transport (Fig. 3B, Table S1, Fig. S4). In contrast, Control Non-Aggressive queens (CNA) exhibited unique GBM genes related to mitochondrial function, oxidative processes, and signal transduction (Fig. 3B, Table S2, Fig. S5). Treated queens (TA and TNA) also had unique genes linked to mitochondrial activity, oxidation processes, and sodium-potassium exchange (Table S4, Fig. S6).

We next examined the methylation profiles of **DNMT1 and DNMT3**, the enzymes responsible for maintaining and de novo DNA methylation. Treated queens (TA and TNA) exhibited lower DNMT1 methylation frequencies compared to control queens (CA and CNA), with the highest methylation observed in CA individuals (Fig. S7). This reduction in DNMT1 methylation in treated individuals is consistent with the mechanism of 5-aza-dC, which incorporates into replicating DNA and irreversibly binds DNMT1, inhibiting maintenance methylation and leading to passive demethylation across cell divisions.

A decrease in DNMT1 methylation—and potentially its expression—may in turn affect the methylation status of downstream target genes, as reflected in the overall weighted methylation levels across gene regions, which were higher in control queens compared to treated ones. Similarly, **DNMT3**, the de novo DNA methyltransferase, also showed lower average methylation levels in treated queens (Fig. S7).

Differential methylation analysis identified 23 genes with significant methylation differences between control aggressive and control non-aggressive queens (Fig. 4, Table S5, Fig S8). These differentially methylated genes were associated with energy metabolism and protein degradation (Fig. 4, Table S4). Additionally, differential methylation analysis identified 334 genes with significant methylation changes between control (CA and CNA) and treated queens (TA and TNA) (Fig. 4, Table S6, Fig S9). Of these, 327 genes showed reduced methylation frequencies in at least 30% of the treated queens (TA and TNA) (Fig. 4, Appendix 4). These genes were primarily associated with housekeeping functions such as RNA and DNA synthesis, peptidyl−arginine methylation, ATP production and mitochondrial functions (e.g., energy metabolism), DNA repair, and general metabolism (Fig. 4, Fig. S10, Appendix 4).

**Figure 4.**
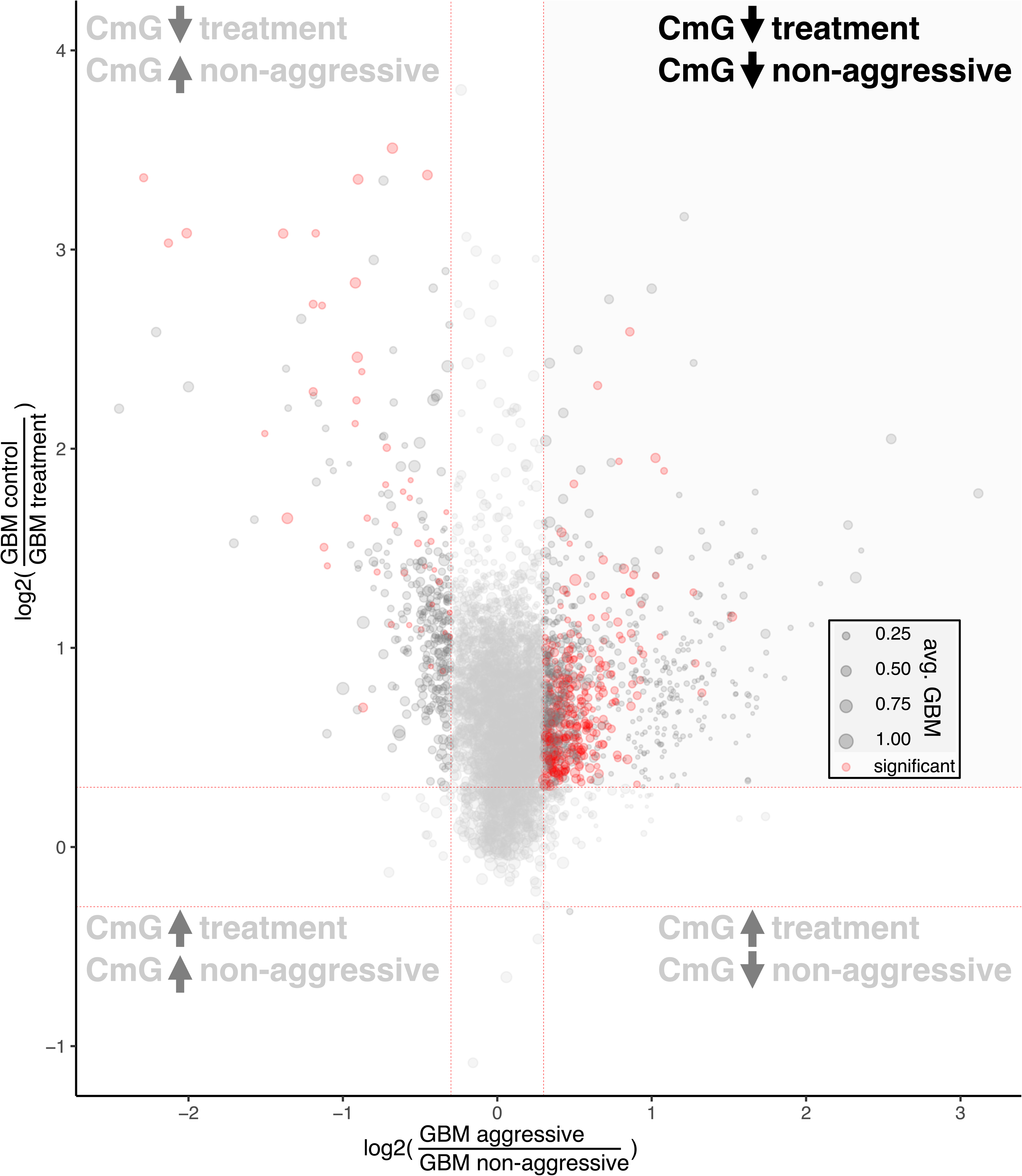
Volcano plot comparing differential expression across two experimental contrasts by plotting the negative log2 fold change from the control aggressive vs. control non-aggressive comparison on the x-axis and the negative log2 fold change from the control vs. treatment comparison on the y-axis. Each point represents a gene, with point size scaled by its average expression across control and treatment, and color indicating whether the gene was classified as significant (red, n= 355 genes) or not (black). Differentially methylated genes were identified by Wilcoxon rank-sum test and thresholds for inclusion were log2-fold change (>0.3=differential methylated; <−0.3=non differential methylated) and Benjamini–Hochberg adjusted (p < 0.05). Dashed vertical and horizontal lines at zero mark the baseline of no fold change, helping to visualize direction and magnitude of expression differences. Tthe y-axis is restricted to the range -0.2 to 1 to focus on relevant variation.

Spearman correlation analysis of differentially methylated genes across queens belonging to all treatments identified 144 genes significantly associated with queen aggression rank (see methods for more details). Of these, the majority (130 genes) showed a positive correlation with aggression (Appendix 1–2). These genes were primarily linked to translation, general metabolism, ATP production, and mitochondrial function, indicating a strong association with energy metabolism (Appendices 2–4; Figs. S10–S11).

### Methylation frequencies comparing delay vs non-delay

Average methylation for gene regions was a little higher (∼0.038%) in the non-delay queens (W = 1498, p-value = 1.802e-05) by a Hodges–Lehmann median difference of 0.00038 (95% CI 0.00022–0.00055; V = 1498; p = 1.8×10⁻□). Rank-biserial r = 0.64 (Fig S12). Differential methylation analysis did not identify any gene that changes methylation levels between delay and non-delay queens (Fig. S13).

## Discussion

The evolution of social polymorphism in *Pogonomyrmex californicus* is thought to enhance colony fitness in response to specific environmental conditions (Boomsma et al. 2014; Saltz et al. 2016). Colonies can develop as either monogynous (haplometrotic) or polygynous (pleometrotic) multifamily units, both of which operate cooperatively as a single colony (Cahan & Julian 1999; Overson et al. 2014). In *P. californicus*, these distinct social forms arise during the nest-founding stage and allopatric populations are almost fixed for one or the other social type (Overson et al. 2014; Erbii et al. 2024). Erbii et al. (2024) showed that two allopatric populations nearly fixed for alternative behavioral phenotypes showed strong genetic differentiations for specific genomic regions (Erbii et al. 2024). Interestingly, a previous study showed that gene expression was only moderately differentiated between both social types (Helmkampf et al.2016). However, there was a strong transcriptional difference between aggressive and non-aggressive queens (Helmkampf et al. 2016). Since both queen types can show aggression the key difference between the haplometrotic and pleometrotic queens is the probability to attack other queens, i.e. initiate aggression. We wanted to investigate whether an epigenetic mechanism like DNA methylation can contribute to these behavioral differences. To explore this, we conducted the first experimental manipulation of genome-wide DNA methylation in ants using the demethylating agent 5-aza-2′- deoxycytidine (5-aza-dC). Importantly, our study employed a decitabine concentration previously used in experiments with bumblebees and *Nasonia* wasps, which reported no deleterious effects on insect health or viability (Cook et al. 2015; Cook et al. 2019; Pozo et al. 2021).

Chemically induced DNA demethylation has been shown to contribute to phenotypic variation in insects (Cook et al. 2015; Cook et al. 2019; Pozo et al. 2021). In this study, we used *P. californicus* founding queens to examine how altered methylation influences behavioral patterns during colony founding, with a focus on aggressive behavior. Our findings revealed that queens exposed to continuous decitabine treatment for 10 days exhibited significantly lower aggression levels compared to untreated control queens.

Methylation profiles between treated and control groups showed distinct differences in methylation frequencies, confirming the genome-wide impact of the DNA methylation inhibitor decitabine. Treated queens (TA and TNA) displayed overall reduced DNA methylation levels, including the two DNA methyltransferases DNMT1 and DNMT3 (Fig. S7). Given that we found no genetic structure between queens groups which differ in aggression and treatment, and founding queens stem from the same population (Fig S3), we infer that the DNA methylation differences were determined by the exposed to the chemical. As expected, most differentially methylated genes (DMGs) in treated queens showed reduced methylation frequencies, with decreases of 26%. Gene ontology analysis revealed that these DMGs were associated with essential biological processes, including RNA and DNA synthesis (housekeeping functions), peptidyl−arginine methylation, ATP production, mitochondrial function, DNA repair, and general metabolism.

These findings are consistent with previous research in *P. californicus*, which reported gene expression differences between aggressive and non-aggressive founding queens. In particular, aggressive queens exhibited upregulation of genes involved in metabolic pathways such as oxidative phosphorylation, including cytochrome c oxidase II and NADH dehydrogenase, both of which are key contributors to ATP production in eukaryotic cells (Anholt & Mackay 2012; Li-Byarlay et al. 2014; Toth et al. 2014; Helmkampf et al. 2016). This pattern aligns with broader evidence across the animal kingdom showing that changes in the expression of metabolic genes are commonly associated with aggressive behavior (Gammie et al. 2007; Alaux et al. 2009; Ayroles et al. 2009; Edwards et al. 2009; Anholt & Mackay 2012; Li-Byarlay et al. 2014; Toth et al. 2014; Helmkampf et al. 2016). Additionally, elevated expression of metabolism-related genes has been observed in the brains of maternally aggressive mice (Gammie et al. 2007) and in the whole bodies of *Drosophila melanogaster* selectively bred for aggressive behavior. Furthermore, bumblebee workers exposed to decitabine exhibited more cooperative behavior, displaying reduced aggresion levels compared to untreated individuals (Pozo et al. 2021).

It has been shown that DNA methylation in insects does not reduce expression as in mammals or plants but rather is linked to higher or constitutional expression. We have shown indirectly that this is also the case in *P. californicus* (Chavarria-Pizarro 2025). Both DNMT1 and DNMT3 were significantly more methylated in untreated founding queens of *P. californicus* compared to treated queens (Fig. S7). As DNMT plays a key role in maintaining DNA methylation during insect development and is involved in regulating gene expression related to the division of labor between queens and workers in social insects such as honeybees (Kucharski et al. 2008; Lyko et al. 2010; Liebrecht et al. 2016) reduction in DNA methylation of these two key genes for DNA methylation should result in lower genome wide DNA methylation. Previous studies have shown that reduced DNMT1 expression leads to demethylation of specific genes (Leu et al. 2003, Suzuki et al. 2004, Bae et al. 2016). Given that genetic variation can influence both DNA methylation and gene expression, future studies should consider genetic background when investigating links between DNA methylation and social polymorphism (Wedd et al. 2016; Errbii et al. 2024).

## Conclusion

This study provides experimental evidence that DNA methylation influences queen behavior during nest founding in *Pogonomyrmex californicus*. By manipulating genome-wide methylation with decitabine, we found that treated queens exhibited significantly reduced aggression compared to controls. Methylation profiling confirmed global hypomethylation in treated individuals, including in genes encoding DNMT1 and DNMT3. Notably, aggressive control queens also showed increased methylation in genes related to energy metabolism, consistent with known links between metabolic gene expression and aggression. These findings support a potential epigenetic basis for behavioral plasticity and social variation in this species’ social niche polymorphism. Future studies using gene-specific methylation knock-downs are needed to establish causal relationships between CpG methylation and behavioral phenotypes.

## Acknowledgments

Hilde Schwitte for performing all the DNA extractions. Kathrin Brüggemann for generating the MinION libraries and running the sequencing. Dr. Jennifer Fewell for hosting our project at the University of Arizona State. Dr. Aline Muyle for generating the R scripts for some methylation analysis used in this manuscript. This work was funded by the German Research Foundation (DFG) as part of the CRC 438 TRR 212 (NC3, 316099922), project C04 (granted to JG).

## Conflict of interested

None declared.

## Author contributions

M.E. contributed by generating the genome annotation (Errbii et al. unpublished), data analysis and manuscript editing. L.S. contributed to data analysis and manuscript editing. B.K contributed to data analysis. M.M assisted to collect the founding queens and the behaviour data for the first experiment season 2023 and for the second one 2024. J.G. conceived the project proposal to obtain the grant, contributed to manuscript writing and editing. T.Ch.P. collected the founding queens for the experiment, designed and performed the experiment, analysed most of the data and wrote the manuscript.

## Data availability statement

The methylation calls have been deposited with links to https://figshare.com/s/a54d6faf3e8687c5483a. The data will be available after peer review publication. The required scripts to do the methylation analysis (e.g. basecalling, methylation calling, bedtools, samtools, and R script) are available in this repositor https://github.com/TaniaChP79/P.cal-methylome

## Notes

https://figshare.com/s/a54d6faf3e8687c5483a

### Competing Interest Statement

The authors have declared no competing interest.

### Summary of Updates

The revision summary has an improve of the figures

## References

Alaux C, S Sinha, L Hasadsri L et al. 2009. Honey bee aggression supports a link between gene regulation and behavioral evolution. Proceedings of the National Academy of Sciences of the USA. 106, 15400–15405. 10.1073/pnas.0907043106

Alexa A & J Rahnenfuhrer. 2024. topGO: Enrichment Analysis for Gene Ontology. R package version 2.56.0.

Amarasinghe HE, CI Clayton, EB Mallon. 2014. Methylation and worker reproduction in the bumble-bee (Bombus terrestris). Proceedings Royal Society B Biology Sciences. 281:20132502. 10.1098/rspb.2013.2502

Anholt RRH & TFC Mackay TFC. 2012. Genetics of aggression Annual Review of Genetics. 46, 145–164. 10.1146/annurev-genet-110711-155514

Ayroles JF, MA Carbone, EA Stone et al. 2009. Systems genetics of complex traits in Drosophila melanogaster. Nature Genetics. 41, 299–307. 10.1038/ng.332

Bernasconi G & J Strassmann. 1999.Cooperation among unrelated individuals: the ant foundress case. Trends in Ecology and Evolution.14, 477–482. doi: 10.1016/S0169-5347(99)01722-X

Berger S L. 2007. The complex language of chromatin regulation during transcription. Nature. 44: 407–412. 10.1038/nature05915

Benjamin Y & Y Hochberg. 1995. Controlling the false discovery rate: a practical and powerful approach to multiple testing. Journal of the Royal Statistical Society (Series B). 57:289–300. 10.1111/j.2517-6161.1995.tb02031.x

Bewick AJ, L Ji, CE Niederhuth et al. 2016. On the origin and evolutionary consequences of gene body DNA methylation. Proceedings of National Academy of Science.113:9111–6. 10.1073/pnas.1604666113

Bewick AJ, JK Vogel, J Allen et al. 2017. Evolution of DNA Methylation across Insects. Molecular Biology and Evolution.34(3): 654–665. 10.1093/molbev/msw264

Bewick AJ, Z Sanchez, EC. Mckinney et al. 2019. Dnmt1 is essential for egg production and embryo viability in the large milkweed bug, Oncopeltus fasciatus. Epigenetics Chromatin. 12:6. 10.1186/s13072-018-0246-5

Bird A. 2007. Perceptions of epigenetics. Nature. 447, 396–398. 10.1038/nature05913

Bolger A M, M Lohse & B Usadel. 2014. Trimmomatic: a flexible trimmer for Illumina sequence data. Bioinformatics. 30(15):2114–20. doi: 10.1093/bioinformatics/btu170. PMID: 24695404; PMCID: PMC4103590.

Bonasio R, Q Li, J Lian et al. 2012. Genome-wide and caste-specific DNA methylomes of the ants Camponotus floridanus and Harpegnathos saltator. Current Biology. 22(19), 1755–1764. doi: 10.1016/j.cub.2012.07.042

Boomsma JJ, DB Huszr & JS Pedersen. 2014. The evolution of multiqueen breeding in eusocial lineages with permanent physically differentiated castes. Animal Behavior. 92:241–52. 10.1016/j.anbehav.2014.03.005.

Cahan S & G Julian.1999. Fitness consequences of cooperative colony founding in the desert leaf-cutter ant Acromyrmex versicolor. Behavioral Ecology. 10, 585–591. 10.1093/beheco/10.5.585

Cardoso-Júnior CAM, B Yagound, I Ronai et al. 2021. DNA methylation is not a driverof gene expression reprogramming in young honey bee workers. Molecular Ecology. 30(19): 4804–4818. 10.1111/mec.16098

Chavarria-Pizarro T, M. Errbii, M. Rinke et al. (2025). Castes and developmental stages of the harvester ant P. californicus differ in genome-wide and gene-specific DNA methylation. bioRxiv. 10.1101/2025.03.13.642958

Cheng X & RM Blumenthal. 2008. Mammalian DNA methyltransferases: a structural perspective. Structure. 16:341–350. doi: 10.1016/j.str.2008.01.004

Clark RM & JH Fewell. 2014. Social dynamics drive selection in cooperative associations of ant queens. Behavioral Ecology. 25, 117–123. 10.1093/beheco/art093

Christman J. K. 2002. 5-Azacytidine and 5-aza-2′-deoxycytidine as inhibitors of DNA methylation: mechanistic studies and their implications for cancer therapy. Oncogene. 21:5483–5495. 10.1038/sj.onc.1205699

Cook, N, B. A. Pannebakker, E. Tauber et al. 2015. DNA methylation and sex allocation in the parasitoid wasp Nasonia vitripennis. American Naturalist. 186:513–518. https://www.journals.uchicago.edu/doi/full/10.1086/682950

Cook N, Parker DJ, F Turner et al. 2019 Genome-wide disruption of DNA methylation by 5-aza-2’deoxycytidine in the parasitoid wasp Nasonia vitripennis. BioRxiv. 2019;437202. doi: 10.1101/437202

Deans C & K A. Maggert What Do You Mean, “Epigenetic”?, Genetics. 199 (4) 887– 896, 10.1534/genetics.114.173492

Edwards AC, Rollmann SM, Morgan TJ et al. 2006. Quantitative genomics of aggressive behavior in Drosophila melanogaster. PLoS Genetics. 2, e154. 10.1371/journal.pgen.0020154

Errbii M. Genome assembly and the raw sequencing data for Pogonomyrmex californicus. 2021. https://www.ncbi.nlm.nih.gov/bioproject/PRJNA682388/. Accessed 25 Apr 2024.

Errbi M, UR Ernst, A Lajmi. et al. 2024. Evolutionary genomics of socially polymorphic populations of Pogonomyrmex californicus. BMC Biology. 22: 109. 10.1186/s12915-024-01907-z

Gammie SC, AP Auger, & HM Jessen et al. 2007. Altered gene expression in mice selected for high maternal aggression. Genes, Brain and Behavior. 6, 432–443. 10.1111/j.1601-183X.2006.00271.x

Grant-Downton R.T. & HG Dickinson. 2006. Epigenetics and its implications for plant biology: 2. The epigenetic epiphany: epigenetics, evolution and beyond. Annals of Botany. 97: 11–27. 10.1093/aob/mcj001

Goll MG & TH Bestor. 2005. Eukaryotic cytosine methyltransferases. Annual Review Biochemestry. 74:481–514. 10.1146/annurev.biochem.74.010904.153721

Hagemann S, O Heil, F. Lyko, et al. 2011. Azacytidine and decitabine induce gene-specific and non-random DNA demethylation in human cancer cell lines. PLOS ONE. 6:e17388. 10.1371/journal.pone.0017388

Helmkampf M, AS Mikheyev, Y Kang, et al. 2016. Gene expression and variation in social aggression by queens of the harvester ant Pogonomyrmex californicus. Molecular Ecology. 25: 3716–30. 10.1111/mec.13700

Herb B R, F Wolschin, K.D. Hansen, et al. 2012. Reversible switching between epigenetic states in honeybee behavioral subcastes. Nature Neuroscience. 15(10): 1371–1373. 10.1038/nn.3218

Holbrook CT, R Clark, R Jeanson et al. 2009. Emergence and consequences of division of labor in forced associations of normally solitary sweat bees. Ethology. 115, 301–310. 10.1111/j.1439-0310.2009.01617.x

Holliday R & JE Pugh. 1975. DNA modification mechanisms and gene activity during development. Science. 187:226–232. DOI: 10.1126/science.187.4173.226

Jablonka E & MJ Lamb. 2002. The changing concept of epigenetics. Annals of the New York Academy of Science. 981:82–96. 10.1111/j.1749-6632.2002.tb04913.x

Johnson R A. 2004. Colony founding by pleometrosis in the semiclaustral seed-harvester ant *Pogonomyrmex californicus* (Hymenoptera: Formicidae). Animal Behaviour. 68:1189–200. 10.1016/j.anbehav.2003.11.021

Kaiser M. I, J Gadau, S Kaiser et al. 2004. Individualized social niches in animals: Theoretical clarifications and processes of niche change. BioScience. 74 (3): 146–158. 10.1093/biosci/biad122)),

Keller L. 1993. The assessment of reproductive success of queens in ants andother social insects. Oikos. 67:177. 10.2307/3545107

Kim JK, M Samaranayake & S Pradhan 2008. Epigenetic mechanisms in mammals. Cellular Molecular Life Science. 66:596–612. 10.1007/s00018-008-8432-4

Kucharski R, J. Maleszka, S Foret & R Maleszka. 2008. Nutritional control of reproductive status in honeybees via DNA methylation. Science. 319(5871): 1827–1830. DOI: 10.1126/science.1153069

Li H. 2018. Minimap2: pairwise alignment for nucleotide sequences. Bioinformatics. 34:3094–100. 10.1093/bioinformatics/bty191

Libbrecht R, P R. Oxley, L Keller & D J. Kronauer. 2016. Robust DNA methylation in the clonal raider ant brain. Current Biology. 26(3), 391–395. doi: 10.1016/j.cub.2015.12.040

Li-Byarlay H, CC Rittschof, JH Massey, et al. 2014. Socially responsive effects of brain oxidative metabolism on aggression. Proceedings of the National Academy of Sciences of the USA. 111, 12533–12537. 10.1073/pnas.141230611

Lyko F. 2018. The DNA methyltransferase family: a versatile toolkit for epigenetic regulation. Nature Review Genetics. 19:81–92. 10.1038/nrg.2017.80

Momparler, R. L. 2005. Epigenetic therapy of cancer with 5-aza-2′-deoxycytidine (decitabine). Seminars in Oncology. 32:443–451. 10.1053/j.seminoncol.2005.07.008.

Mossman D, K Kim & RJ Scott. 2010. Demethylation by 5-aza-2′-deoxycytidine in colorectal cancer cells targets genomic DNA whilst promoter CpG island methylation persists. BMC Cancer. 10:366. 10.1186/1471-2407-10-366

Muyle A, J Ross-Ibarra, DK Seymour et al. 2021. Gene body methylation is under selection in Arabidopsis thaliana. Genetics. 218:iyab061. 10.1093/genetics/iyab061

Overson R, J Gadau, RM Clark et al. 2014. Behavioral transitions with the evolution of cooperative nest founding by harvester ant queens. Behavioral Ecology and Sociobiology. 68, 21–30. 10.1007/s00265-013-1618-2

Overson R, JH Fewell & J Gadau. 2016. Distribution and origin of intraspecific social variation in the California harvester ant Pogonomyrmex californicus. Insects Society.63:531–41. 10.1007/s00040-016-0497-8

Patalano S, A Vlasova, C Wyatt et al. 2015. Molecular signatures of plastic phenotypes in two eusocial insect species with simple societies. Proceedings of the National Academy of Sciences of the United States of America.112(45), 13970– 13975. 10.1073/pnas.1515937112

Perez M, O Aroh, Y Sun, Y Lan, et al. 2023. Third-Generation Sequencing Reveals the Adaptive Role of the Epigenome in Three Deep-Sea Polychaetes. Molecular Biology and Evolution. 40 (8): msad172, 10.1093/molbev/msad172

Pozo MI, J Benjamin J. Hunt, G Van Kemenade., et al. 2021. The effect of DNA methylation on bumblebee colony development. BMC Genomics. 22:73 10.1186/s12864-021-07371-1

Purcell J, A Brelsford, Y Wurm et al. 2014.Convergent genetic architecture underlies social organization in ants. Current Biology. 24:2728–32. doi: 10.1016/j.cub.2014.09.071

Quinlan A R, I M Hall. 2010. BEDTools: a flexible suite of utilities for comparing genomic features, Bioinformatics. 26, 6, 841–842. 10.1093/bioinformatics/btq033

Ramos M, NA Wijetunga, AS McLellan et al. 2015. DNA demethylation by 5-aza-2′- deoxycytidine is imprinted, targeted to euchromatin and has limited transcriptional consequences. Epigenetics and Chromatin. 8:11. 10.1186/s13072-015-0004-x

Richards EJ.2006. Inherited epigenetic variation – revisiting soft inheritance. Nature Review Genetics. 7. 395–401. 10.1038/nrg1834

Rissing SW, RA Johnson & JW Martin. 2000. Colony founding behavior of some desert ants: geographic variation in metrosis. Psyche: A Journal of Entomology.103:95–101. 10.1155/2000/20135

Saltz JB, AP Geiger, R Anderson, et al. 2016. What, if anything,is a social niche? Evolutionary Ecology. 30:349–64. 10.1007/s10682-015-9792-5

Seelan RS, P Mukhopadhyay, MM Pisano et al. 2018. Effects of 5-Aza-2′-deoxycytidine (decitabine) on gene expression. Drug Metabolism Reviews. 50:193–207. 10.1080/03602532.2018.1437446

Simpson JT, RE Workman, PC Zuzarte et al. 2017. Detecting DNA cytosine methylation using nanopore sequencing. Nature Methods. 14:407–410. 10.1038/nmeth.4184

Standage DS, AJ Berens, KM.Glastad, et al. 2016. Genome, transcriptome and methylome sequencing of a primitively eusocial wasp reveal a greatly reduced DNA methylation system in a social insect. Molecular Ecology. 25:1769–1784. 10.1111/mec.13578

Takuno S & BS. Gaut. 2012. Body-methylated genes in Arabidopsis thaliana are functionally important and evolve slowly. Molecular Biology Evolution. 29:219–27. 10.1093/molbev/msr188

Toth AL, JF Tooker, S Radhakrishnan et al. 2014. Shared genes related to aggression, rather than chemical communication, are associated with reproductive dominance in paper wasps (Polistes metricus). BMC Genomics. 15, 75. 10.1186/1471-2164-15-75

Wang J, Y Wurm, M Nipitwattanaphon, et al. 2013. A Y-like social chromosome causes alternative colony organization in fire ants. Nature. 493:664–8. 10.1038/nature11832

Wedd L, R Kucharski & R Maleszka. 2016. Differentially methylated obligatory epialleles modulate context-dependent LAM gene expression in the honeybee Apis mellifera. Epigenetics. 11(1):1–10. doi: 10.1080/15592294.2015.1107695. PMID: 26507253; PMCID: PMC4846127

Wong AH, Gottesman II & A Petronis. 2005. Phenotypic differences in genetically identical organisms: the epigenetic perspective. Human Molecular Genetic. 14:R11–8. 10.1093/hmg/ddi116

Yan H, R Bonasio, DF Simola et al.. 2015. DNA methylation in social insects : How epigenetics can control behavior and longevity. Annual Review of Entomology. 60(60): 435–452. 10.1146/annurev-ento-010814-020803

Yang S, C Guo, X Zhao et al. 2017. Divergent methylation pattern in adult stage between two forms of Tetranychus urticae (Acari: Tetranychidae). Insect Science. 25: 667–678. 10.1111/1744-7917.12444

