## Supplementary material for "Epigenetic reprogramming by 5-aza-dC alters gene methylation and suppresses aggression in founding queens of the California harvester ant *Pogonomyrmex californicus*": Supplementary Info Demethylation paper copy.docx

This file includes:

Figs. S1- S13

Tables S1-S4

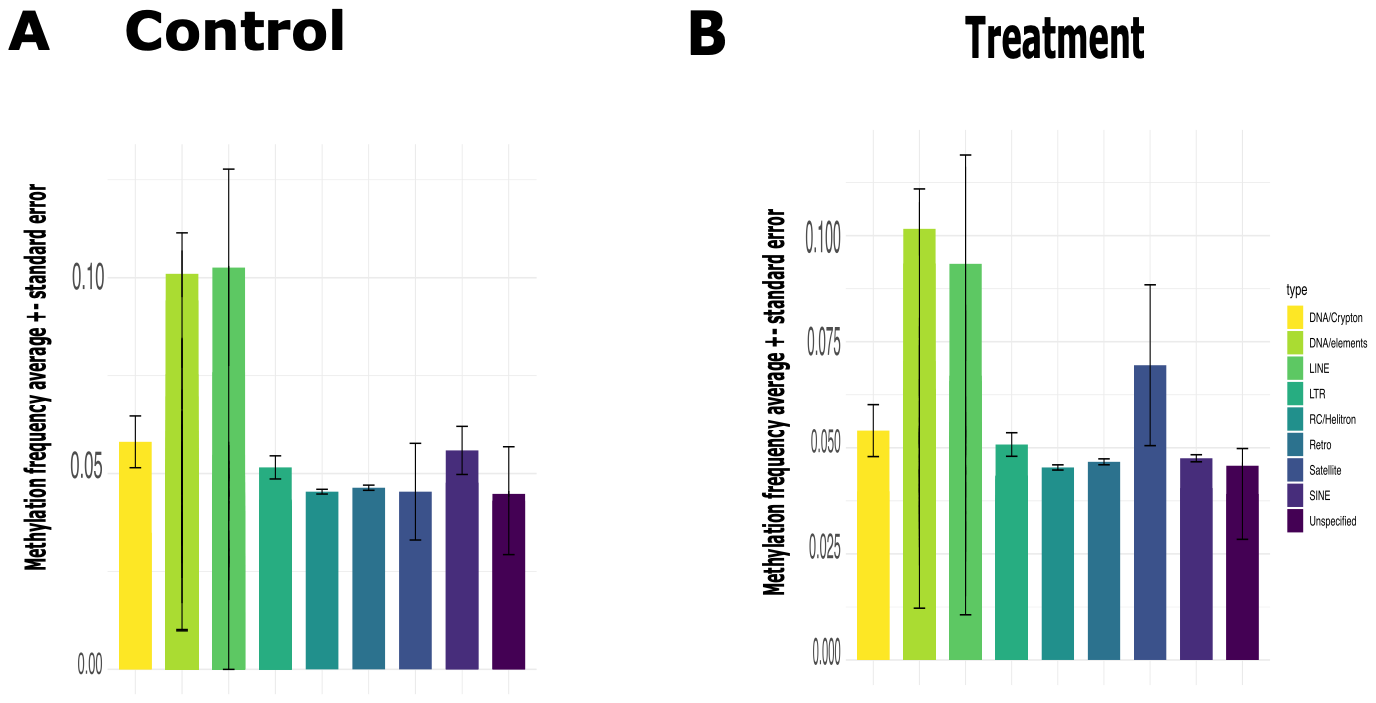

Figure S1. Average Methylation frequency and standard deviation of Transposable Elements (TEs) families in *P. californicus* queen genomes treated (n=12) vs untreated (control)(n=12).

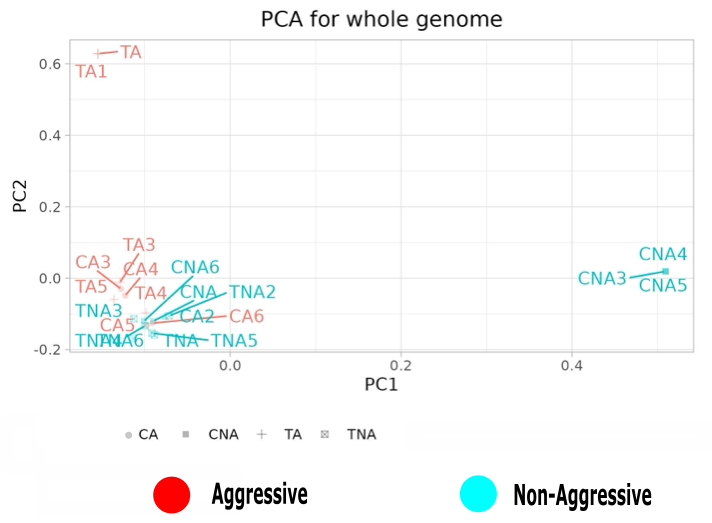

Figure S2. Principal component analysis of genotypes (SNPs) for founding queens

classified individuals into four categories: Control aggressive (CA) (blue circle), Control non-aggressive (CNA) (blue square), Treatment aggressive (TA)(red cross), Treatment non-aggressive (TNA)(red square cross)(n=20).

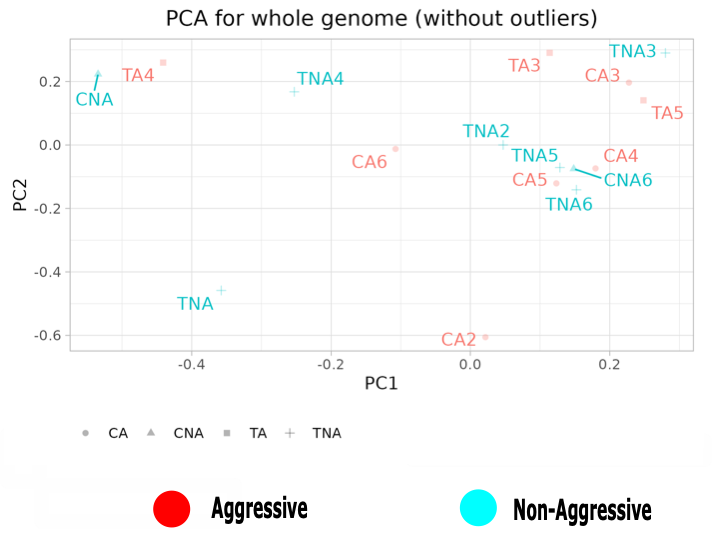

Figure S3. Principal component analysis of genotype (SNPs) without outliers for founding queens classified individuals into four categories: Control aggressive (CA) (blue circle), Control non-aggressive (CNA) (blue square), Treatment aggressive (TA)(red cross), Treatment non-aggressive (TNA)(red square cross)(n=20).

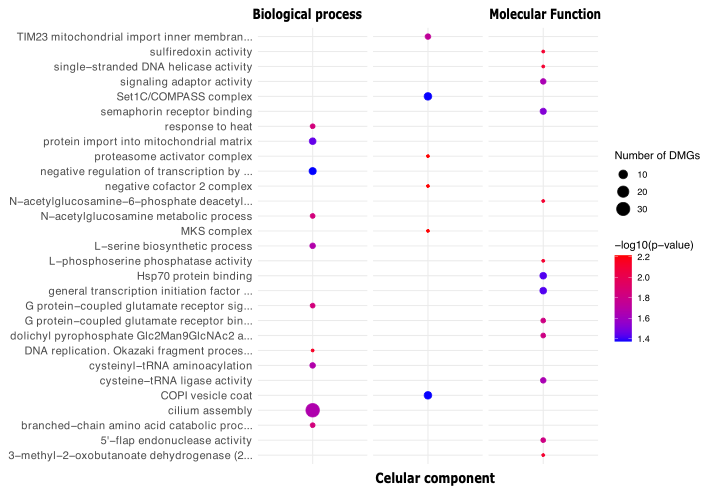

Figure S4. GO enrichment analysis for all genes body methylated unique to queen untreaded (control) aggressive(n=6) with significant gene-body methylation with a p-value < 0.05 representing significantly enriched GO terms.

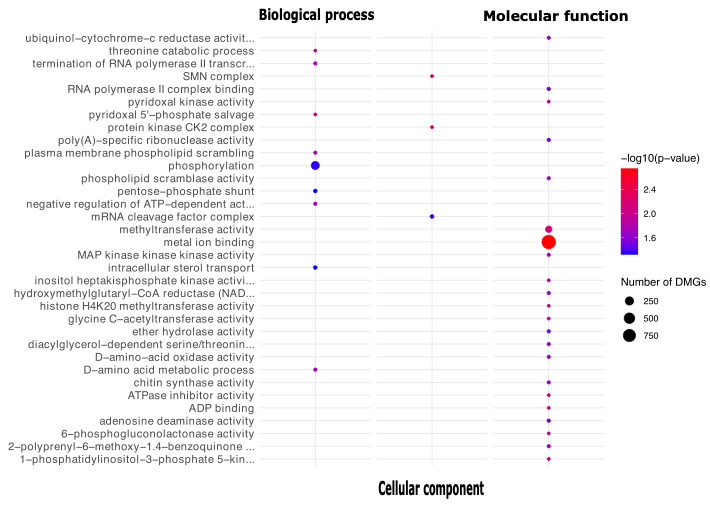

Figure S5. GO enrichment analysis for all genes body methylated unique to queens untreaded (control) non aggressive (n=6) with significant gene-body methylation with a p-value < 0.05 representing significantly enriched GO terms.

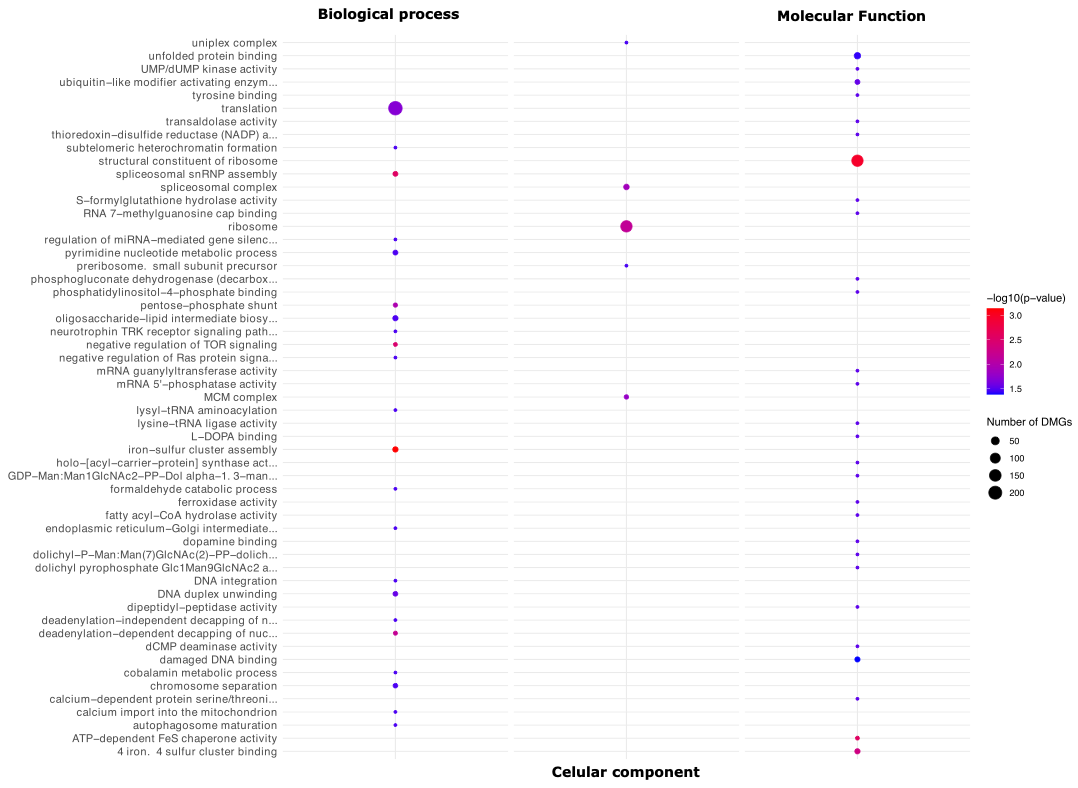

Figure S6. GO enrichment analysis for all genes body methylated unique to queens aza-dC-treated with non aggressive (n=12) with significant gene-body methylation with a p-value < 0.05 representing significantly enriched GO terms.

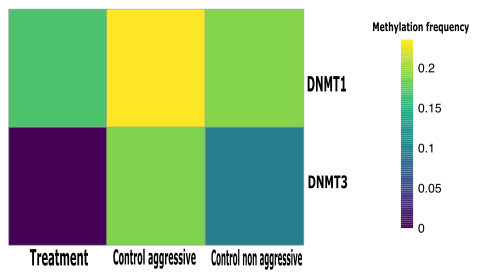

Figure S7. DNMT1 and DNMT3 methylation frequency in *P. californicus* . Average methylation frequency (%) of DNMT1 and DNMT3 along queen aza-dC-treated (n=12) and untreated aggressive (n=6) and untreated non aggressive (n=6).

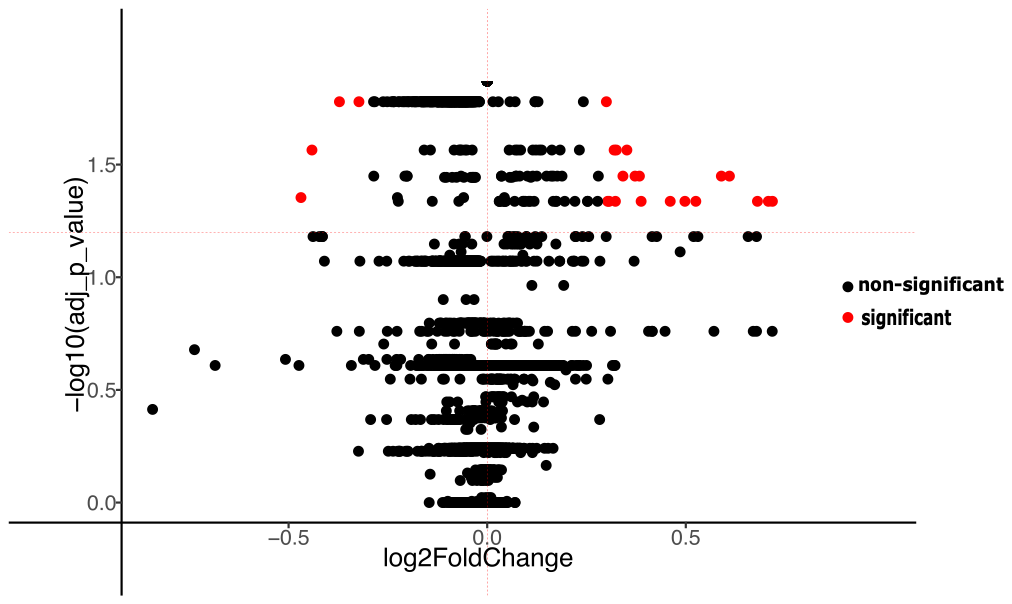

Figure S8. Volcano plot showing differentially methylated genes between control aggressive and control-nonaggressive which were identified by Wilcoxon rank-sum test and thresholds for inclusion were log2-fold change (>0.3=differential methylated; <−0.3=non differential methylated) and Benjamini–Hochberg adjusted (p < 0.05). The p-value < 0.05 in the plot represents significantly red and not significant black.

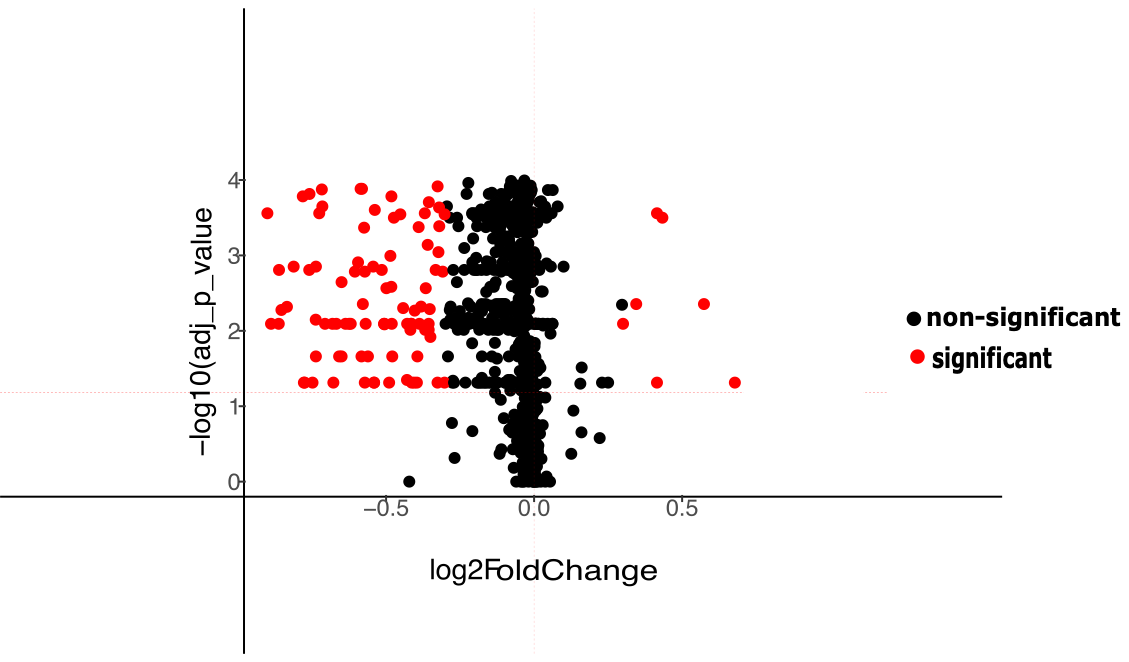

Figure S9. Volcano plot showing differentially methylated genes between control and treatment which were identified by Wilcoxon rank-sum test and thresholds for inclusion were log2-fold change (>0.3=differential methylated; <−0.3=non differential methylated) and Benjamini–Hochberg adjusted (p < 0.05). The p-value < 0.05 in the plot represents significantly red and not significant black.

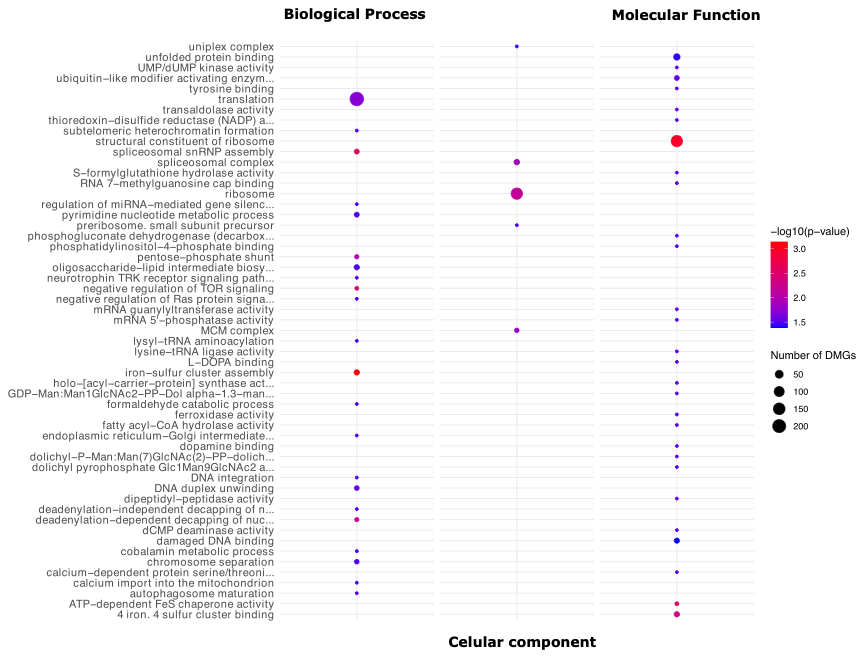

Figure S10. GO enrichment analysis for all genes body methylated which were differentially methylated between comparisons (untreated aggressive vs untreated non aggresiive, untreated vs treated). Differentially methylated genes were identified by Wilcoxon rank-sum test and thresholds for inclusion were log2-fold change (>0.3=differential methylated; <−0.3=non differential methylated) and Benjamini–Hochberg adjusted (p < 0.05). The p-value < 0.05 in the plot represents significantly enriched GO terms.

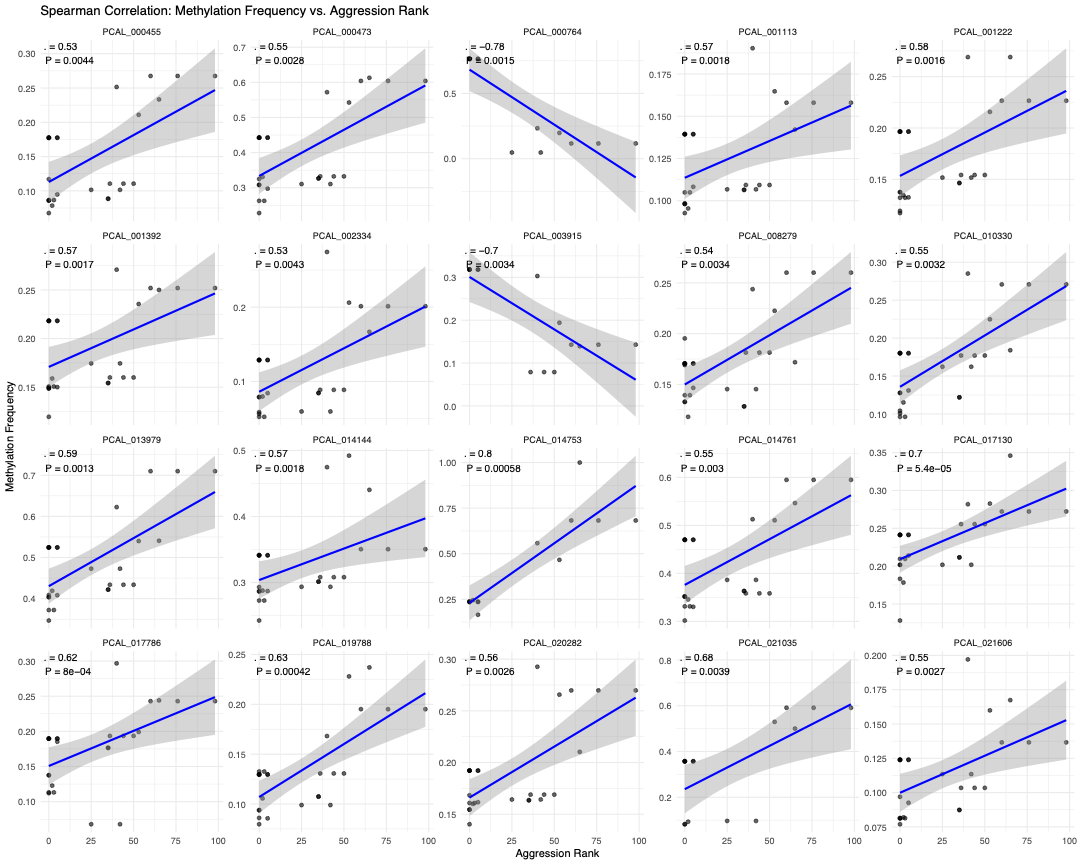

Figure S11. Spearman correlation between queen’s aggression rank (see Behavioral Analysis for details) and the methylation frequencies of the 20 genes that showed higher significant differences in methylation across comparisons (untreated aggressive vs untreated non aggresiive, untreated vs treated).

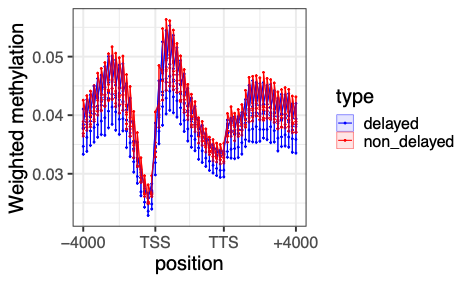

Figure S12. Gene methylation patterns in founding queens along non-delay and delay (control 24) obtain with the sequencing method WGBS A) Weighted frequency methylation average among treatments (n=10 non-delay, n=6 delay) looped over all genes region within a window of 4 kb upstream of the transcription start site (TSS) and a window of 4 kb downstream of the transcription end site (TTS).

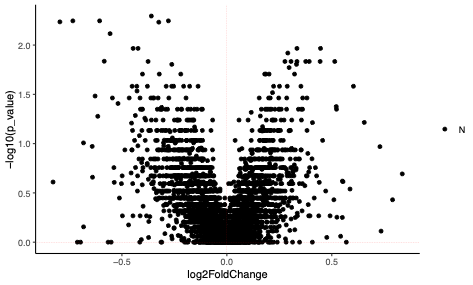

Figure S13. Volcano plot showing differentially methylated genes between delay and non-delay (control 24) which were identified by Wilcoxon rank-sum test and thresholds for inclusion were log2-fold change (>0.3=differential methylated; <−0.3=non differential methylated) and Benjamini–Hochberg adjusted (p < 0.05). The p-value < 0.05 in the plot represents significantly red and not significant black.

Table S1. Gene differential methylated unique for aggressive control queens significant associated to Ontology using a Fisher test. GO.ID (GO Term Identification number), NumDMGs (number differential body methylated genes), pvalue (Fisher test).

| GO.ID | GOTerm | NumDMGs | pvalue | Ontology |
| --- | --- | --- | --- | --- |
| GO:0008448 | N-acetylglucosamine-6-phosphate deacetyl... | 1 | 0.0077 | Molecular_Function |
| GO:0036424 | L-phosphoserine phosphatase activity | 1 | 0.0077 | Molecular_Function |
| GO:0032542 | sulfiredoxin activity | 1 | 0.0077 | Molecular_Function |
| GO:0003863 | 3-methyl-2-oxobutanoate dehydrogenase (2... | 1 | 0.0077 | Molecular_Function |
| GO:0017116 | single-stranded DNA helicase activity | 1 | 0.0077 | Molecular_Function |
| GO:0106073 | dolichyl pyrophosphate Glc2Man9GlcNAc2 a... | 2 | 0.0153 | Molecular_Function |
| GO:0035256 | G protein-coupled glutamate receptor bin... | 2 | 0.0153 | Molecular_Function |
| GO:0017108 | 5'-flap endonuclease activity | 2 | 0.0153 | Molecular_Function |
| GO:0035591 | signaling adaptor activity | 3 | 0.0229 | Molecular_Function |
| GO:0004817 | cysteine-tRNA ligase activity | 3 | 0.0229 | Molecular_Function |
| GO:0030215 | semaphorin receptor binding | 4 | 0.0304 | Molecular_Function |
| GO:0140223 | general transcription initiation factor ... | 5 | 0.0378 | Molecular_Function |
| GO:0030544 | Hsp70 protein binding | 5 | 0.0378 | Molecular_Function |
| GO:0033567 | DNA replication. Okazaki fragment proces... | 1 | 0.0072 | Biological_Process |

Table S1. Continued

| GO.ID | GOTerm | NumDMGs | pvalue | Ontology |
| --- | --- | --- | --- | --- |
| GO:0006044 | N-acetylglucosamine metabolic process | 2 | 0.0143 | Biological_Process |
| GO:0009408 | response to heat | 2 | 0.0143 | Biological_Process |
| GO:0009083 | branched-chain amino acid catabolic proc... | 2 | 0.0143 | Biological_Process |
| GO:0007216 | G protein-coupled glutamate receptor sig... | 2 | 0.0143 | Biological_Process |
| GO:0006423 | cysteinyl-tRNA aminoacylation | 3 | 0.0214 | Biological_Process |
| GO:0006564 | L-serine biosynthetic process | 3 | 0.0214 | Biological_Process |
| GO:0060271 | cilium assembly | 32 | 0.0218 | Biological_Process |
| GO:0030150 | protein import into mitochondrial matrix | 5 | 0.0355 | Biological_Process |
| GO:0000122 | negative regulation of transcription by ... | 6 | 0.0424 | Biological_Process |
| GO:0008537 | proteasome activator complex | 1 | 0.0061 | Cellular_Component |
| GO:0017054 | negative cofactor 2 complex | 1 | 0.0061 | Cellular_Component |
| GO:0036038 | MKS complex | 1 | 0.0061 | Cellular_Component |
| GO:0005744 | TIM23 mitochondrial import inner membran... | 3 | 0.0183 | Cellular_Component |
| GO:0048188 | Set1C/COMPASS complex | 7 | 0.0423 | Cellular_Component |
| GO:0030126 | COPI vesicle coat | 7 | 0.0423 | Cellular_Component |

Table S2. Gene differential methylated unique for non-aggressive control queens significant associated to Ontology using a Fisher test. GO.ID (GO Term Identification number), NumDMGs (number differential body methylated genes), pvalue (Fisher test).

| GO.ID | GOTerm | NumDMGs | pvalue | Ontology |
| --- | --- | --- | --- | --- |
| GO:0046872 | metal ion binding | 981 | 0.0018 | Molecular_Function |
| GO:0008168 | methyltransferase activity | 106 | 0.0067 | Molecular_Function |
| GO:0017057 | 6-phosphogluconolactonase activity | 1 | 0.0107 | Molecular_Function |
| GO:0008890 | glycine C-acetyltransferase activity | 1 | 0.0107 | Molecular_Function |
| GO:0043531 | ADP binding | 1 | 0.0107 | Molecular_Function |
| GO:0042030 | ATPase inhibitor activity | 1 | 0.0107 | Molecular_Function |
| GO:0008478 | pyridoxal kinase activity | 1 | 0.0107 | Molecular_Function |
| GO:0000829 | inositol heptakisphosphate kinase activi... | 1 | 0.0107 | Molecular_Function |
| GO:0042799 | histone H4K20 methyltransferase activity | 1 | 0.0107 | Molecular_Function |
| GO:0000285 | 1-phosphatidylinositol-3-phosphate 5-kin... | 1 | 0.0107 | Molecular_Function |
| GO:0004697 | diacylglycerol-dependent serine/threonin... | 2 | 0.0213 | Molecular_Function |
| GO:0004709 | MAP kinase kinase kinase activity | 2 | 0.0213 | Molecular_Function |
| GO:0008425 | 2-polyprenyl-6-methoxy-1.4-benzoquinone ... | 2 | 0.0213 | Molecular_Function |
| GO:0004100 | chitin synthase activity | 2 | 0.0213 | Molecular_Function |
| GO:0008121 | ubiquinol-cytochrome-c reductase activit... | 2 | 0.0213 | Molecular_Function |

Table S2. Continued.

| GO.ID | GOTerm | NumDMGs | pvalue | Ontology |
| --- | --- | --- | --- | --- |
| GO:0004420 | hydroxymethylglutaryl-CoA reductase (NAD... | 2 | 0.0213 | Molecular_Function |
| GO:0003884 | D-amino-acid oxidase activity | 2 | 0.0213 | Molecular_Function |
| GO:0017128 | phospholipid scramblase activity | 2 | 0.0213 | Molecular_Function |
| GO:0004000 | adenosine deaminase activity | 3 | 0.0318 | Molecular_Function |
| GO:0016803 | ether hydrolase activity | 3 | 0.0318 | Molecular_Function |
| GO:0000993 | RNA polymerase II complex binding | 3 | 0.0318 | Molecular_Function |
| GO:0004535 | poly(A)-specific ribonuclease activity | 3 | 0.0318 | Molecular_Function |
| GO:0006567 | threonine catabolic process | 1 | 0.0097 | Biological_Process |
| GO:0009443 | pyridoxal 5'-phosphate salvage | 1 | 0.0097 | Biological_Process |
| GO:0006369 | termination of RNA polymerase II transcr... | 2 | 0.0192 | Biological_Process |
| GO:0017121 | plasma membrane phospholipid scrambling | 2 | 0.0192 | Biological_Process |
| GO:0046416 | D-amino acid metabolic process | 2 | 0.0192 | Biological_Process |
| GO:0032780 | negative regulation of ATP-dependent act... | 2 | 0.0192 | Biological_Process |
| GO:0016310 | phosphorylation | 235 | 0.0445 | Biological_Process |

Table S2. Continued

| GO.ID | GOTerm | NumDMGs | pvalue | Ontology |
| --- | --- | --- | --- | --- |
| GO:0032366 | intracellular sterol transport | 5 | 0.0474 | Biological_Process |
| GO:0006098 | pentose-phosphate shunt | 5 | 0.0474 | Biological_Process |
| GO:0005956 | protein kinase CK2 complex | 1 | 0.0061 | Cellular_Component |
| GO:0032797 | SMN complex | 1 | 0.0061 | Cellular_Component |
| GO:0005849 | mRNA cleavage factor complex | 7 | 0.0423 | Cellular_Component |

Table S3. Gene differential methylated unique for treated queens significant associated to Ontology using a Fisher test. GO.ID (GO Term Identification number), NumDMGs (number differential body methylated genes), pvalue (Fisher test).

| GO.ID | GOTerm | NumDMGs | pvalue | Ontology |
| --- | --- | --- | --- | --- |
| GO:0004719 | protein-L-isoaspartate (D-aspartate) O-m... | 3 | 5.8e-05 | Molecular_Function |
| GO:0004326 | tetrahydrofolylpolyglutamate synthase ac... | 1 | 0.0045 | Molecular_Function |
| GO:0004767 | sphingomyelin phosphodiesterase activity | 3 | 0.0134 | Molecular_Function |
| GO:0004596 | peptide alpha-N-acetyltransferase activi... | 3 | 0.0134 | Molecular_Function |
| GO:0003682 | chromatin binding | 8 | 0.0354 | Molecular_Function |
| GO:0019888 | protein phosphatase regulator activity | 8 | 0.0354 | Molecular_Function |
| GO:0001736 | establishment of planar polarity | 2 | 0.007 | Biological_Process |
| GO:0006685 | sphingomyelin catabolic process | 2 | 0.007 | Biological_Process |
| GO:0009396 | folic acid-containing compound biosynthe... | 3 | 0.01 | Biological_Process |
| GO:0006479 | protein methylation | 11 | 0.038 | Biological_Process |
| GO:0035145 | exon-exon junction complex | 2 | 0.0049 | Cellular_Component |
| GO:0000159 | protein phosphatase type 2A complex | 3 | 0.0074 | Cellular_Component |
| GO:0005634 | nucleus | 472 | 0.0132 | Cellular_Component |

Table S4. Differential methylated analysis identified 23 genes with significant methylation changes between control aggressive (CA) and control nonaggressive queens (CNA).

| Gene | sites | Genenumber | Geneid | Function | mean_CA | mean_CNA | log2FoldChange | adj_p_value |
| --- | --- | --- | --- | --- | --- | --- | --- | --- |
| PCAL_000769 | Gene3D | G3DSA:1.10.390.10 | Neutral Protease Domain 2 | Peptidase M4/M1, CTD superfamily | 0.1563 | 0.764 | 0.61 | 0.035 |
| PCAL_016166 | Gene3D | G3DSA:2.40.70.10 | Acid Proteases | Aspartic peptidase domain superfamily | 0.528 | 1 | 0.387 | 0.046 |
| PCAL_025664 | Gene3D | G3DSA:1.20.58.2190 | - | - | 0.624 | 1 | 0.3000 | 0.0166 |
| PCAL_004829 | Pfam | PF14937 | Domain of unknown function (DUF4500) | UPF0708 protein C6orf162 | 0.1625 | 0.6 | 0.460 | 0.046 |
| PCAL_010969 | SMART | SM00220 | serkin_6 | Protein kinase domain | 0.513 | 0.958 | 0.372 | 0.0355 |
| PCAL_004141 | CDD | cd00170 | SEC14 | CRAL-TRIO lipid binding domain | 0.35 | 0.667 | 0.303 | 0.046 |
| PCAL_007641 | ProSiteProfiles | PS50031 | EH domain profile. | EH domain | 0.250 | 0.56 | 0.32 | 0.027 |
| PCAL_004141 | CDD | cd00170 | SEC14 | CRAL-TRIO lipid binding domain | 0.35 | 0.667 | 0.303 | 0.046 |
| PCAL_007641 | ProSiteProfiles | PS50031 | EH domain profile. | EH domain | 0.250 | 0.56 | 0.32 | 0.027 |

Table S4. Continued

| Gene | Sites | Genenumber | Geneid | function | mean_CA | mean_CNA | log2FoldChange | adj_p_value |
| --- | --- | --- | --- | --- | --- | --- | --- | --- |
| PCAL_020415 | PANTHER | PTHR36159 | PROTEIN CBG23766 | - | 0.34 | 0.89 | 0.497 | 0.046 |
| PCAL_003823 | Coils | Coil | Coil | - | 0.50 | 0.199 | -0.323 | 0.0167 |
| PCAL_021711 | PANTHER | PTHR12395:SF9 | DECAPPING AND EXORIBONUCLEASE PROTEIN | - | 0.224 | 1 | 0.71 | 0.046 |
| PCAL_026098 | Gene3D | G3DSA:3.15.10.30 | Haemolymph juvenile hormone binding protein | Takeout superfamily | 0.53 | 1 | 0.383 | 0.035 |
| PCAL_010976 | Coils | Coil | Coil | - | 0.13 | 0.411 | 0.32 | 0.046 |
| PCAL_007393 | Pfam | PF06522 | NADH-ubiquinone reductase complex 1 MLRQ subunit | NADH-ubiquinone reductase complex 1 MLRQ subunit | 0.389 | 1 | 0.525 | 0.046 |
| PCAL_001710 | Gene3D | G3DSA:3.30.160.20 | - | - | 0.248 | 1 | 0.68 | 0.046 |
| PCAL_015881 | Gene3D | G3DSA:2.110.10.10 | - | Hemopexin-like domain superfamily | 0.68 | 0.24 | -0.44 | 0.027 |
| PCAL_019026 | Pfam | PF12259 | Baculovirus F protein | Envelope fusion protein-like | 0.273 | 0.625 | 0.3517 | 0.027 |

Table S4. Continued.

| Gene | Sites | Genenumber | Geneid | function | mean_CA | mean_CNA | log2FoldChange | adj_p_value |
| --- | --- | --- | --- | --- | --- | --- | --- | --- |
| PCAL_020731 | MobiDBLite | mobidb-lite | consensus disorder prediction | - | 0.262 | 0.6 | 0.341 | 0.0356 |
| PCAL_001865 | MobiDBLite | mobidb-lite | consensus disorder prediction | - | 0.216 | 1 | 0.718 | 0.046 |
| PCAL_000764 | Gene3D | G3DSA:2.60.40.1910 | - | - | 0.469 | 0.83 | 0.32 | 0.027 |
| PCAL_014753 | Pfam | PF00348 | Polyprenyl synthetase | Polyprenyl synthetase | 0.17 | 0.769 | 0.59 | 0.035 |
| PCAL_012803 | PANTHER | PTHR36159 | PROTEIN CBG23766 | - | 0.809 | 0.397 | -0.372 | 0.0166 |
| PCAL_000764 | Gene3D | G3DSA:1.10.390.10 | Neutral Protease Domain 2 | Peptidase M4/M1, CTD superfamily | 0.906 | 0.377 | -0.468 | 0.044 |
