## Supplementary figures and images for "Epigenetic reprogramming by 5-aza-dC alters gene methylation and suppresses aggression in founding queens of the California harvester ant *Pogonomyrmex californicus*"

### Fig S1.pdf

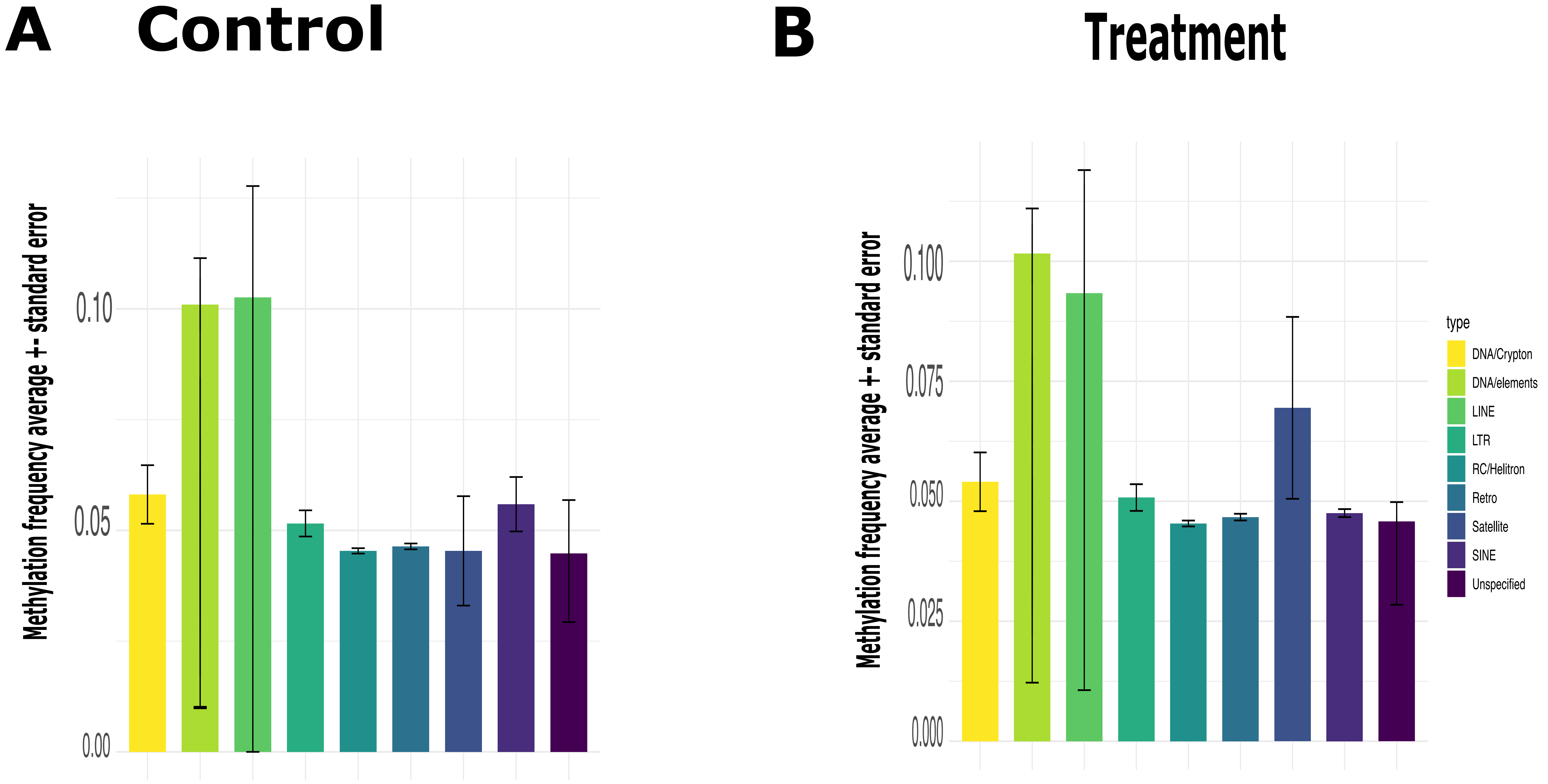

### Fig S2. PCA with outliers. pdf.pdf

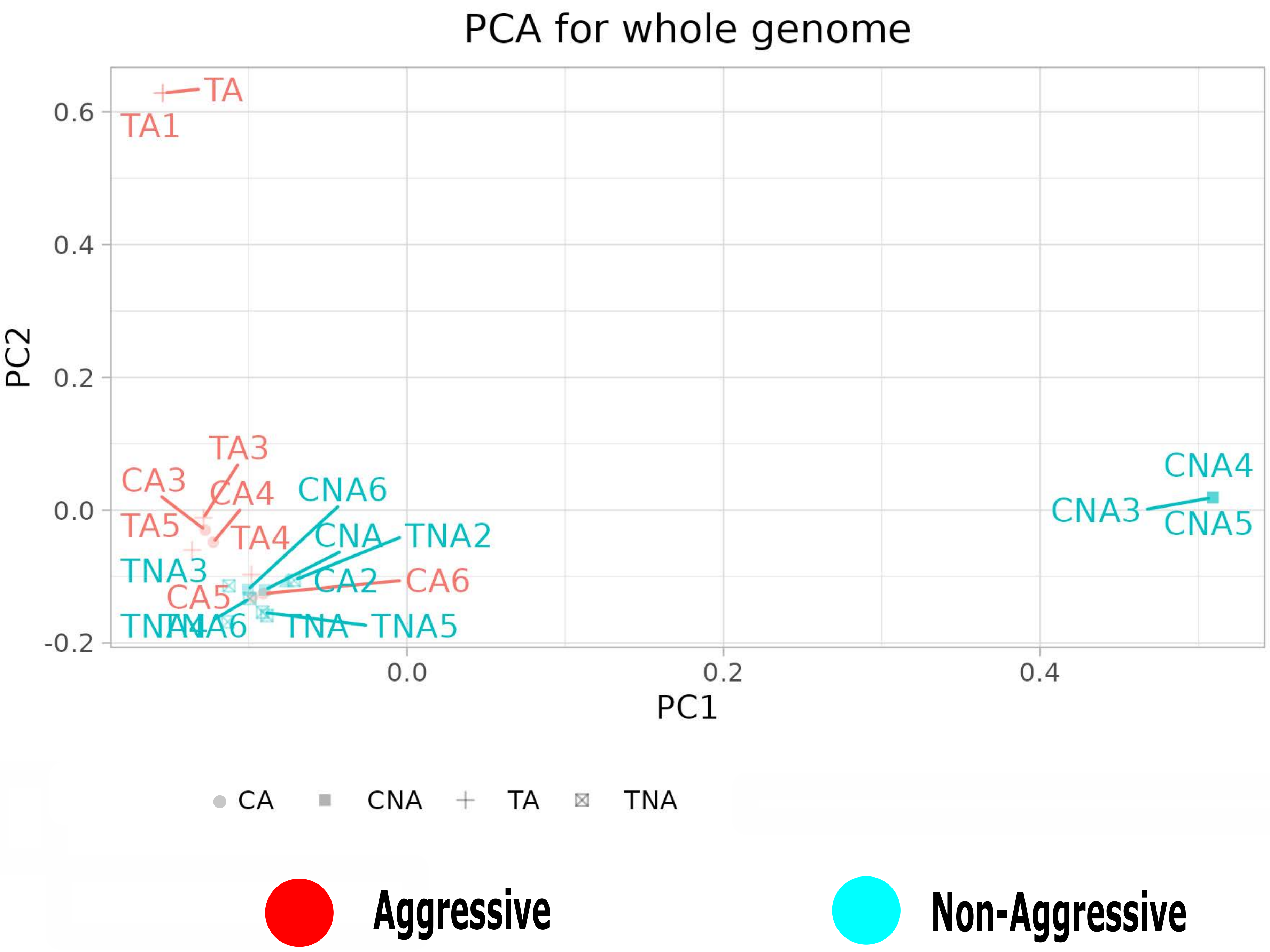

### Fig S3. PCA without outliers.pdf

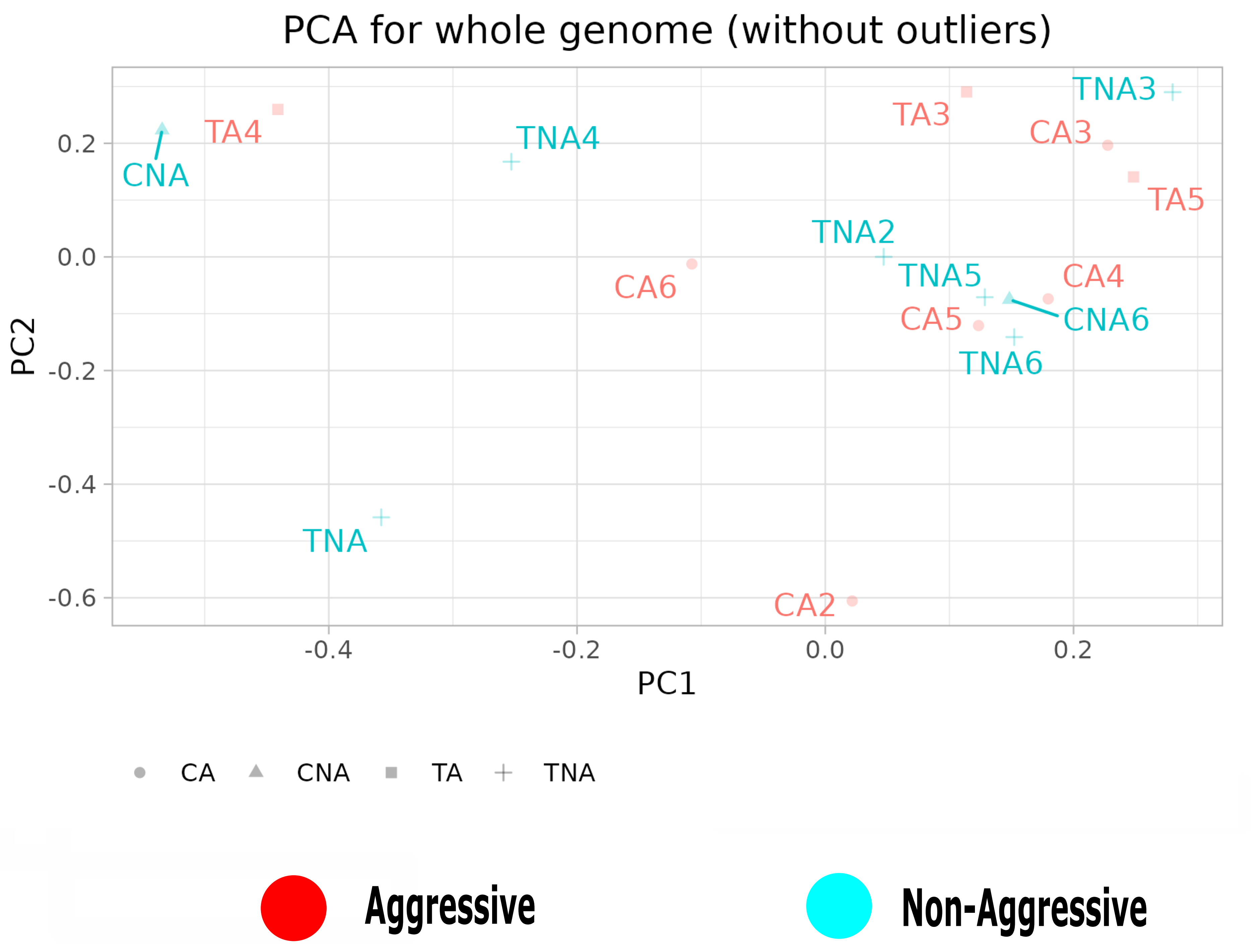

### Fig S4.DMGnewControlaggressive.pdf

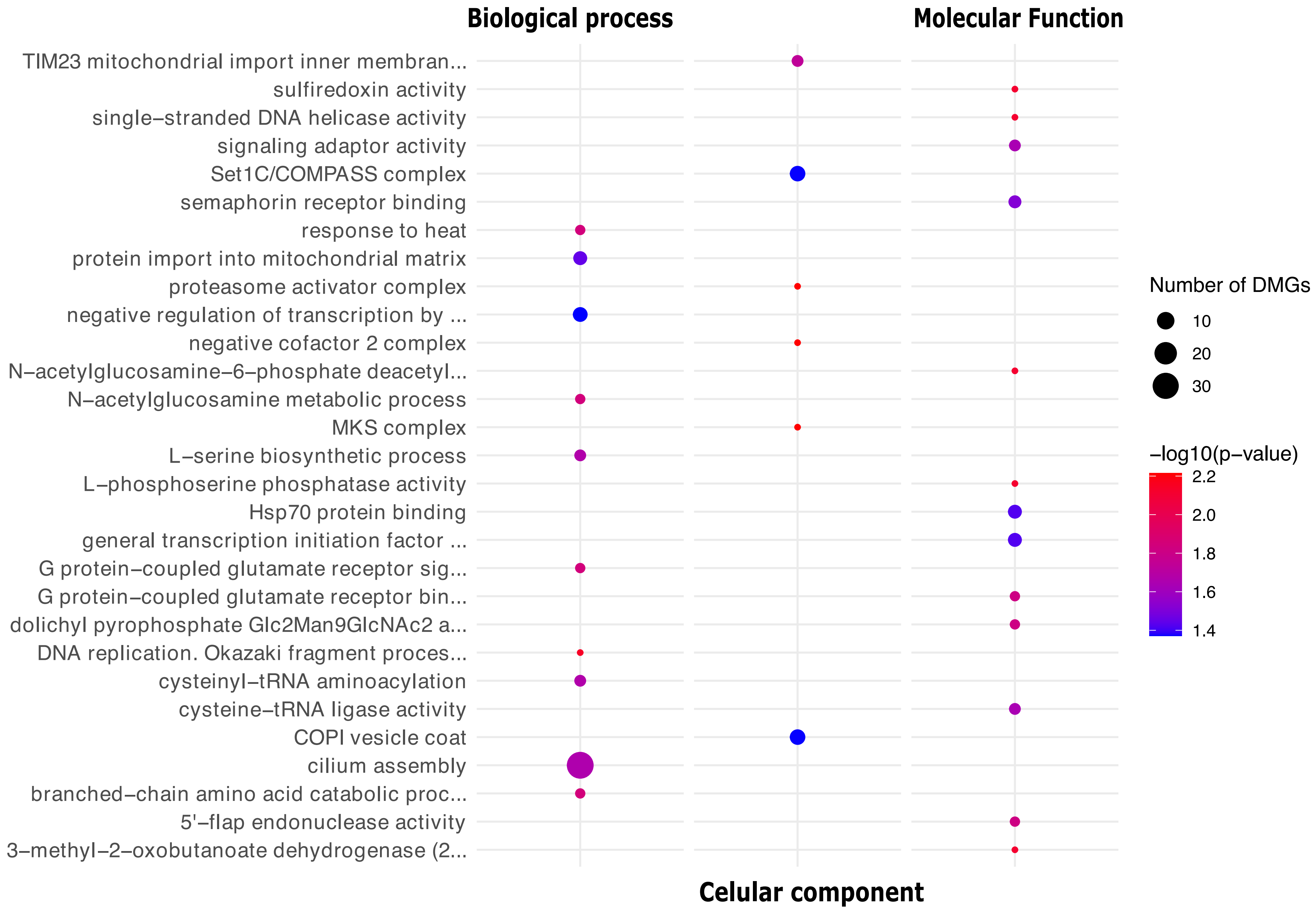

### Fig S5. DMGnewControlNonAggressive.pdf

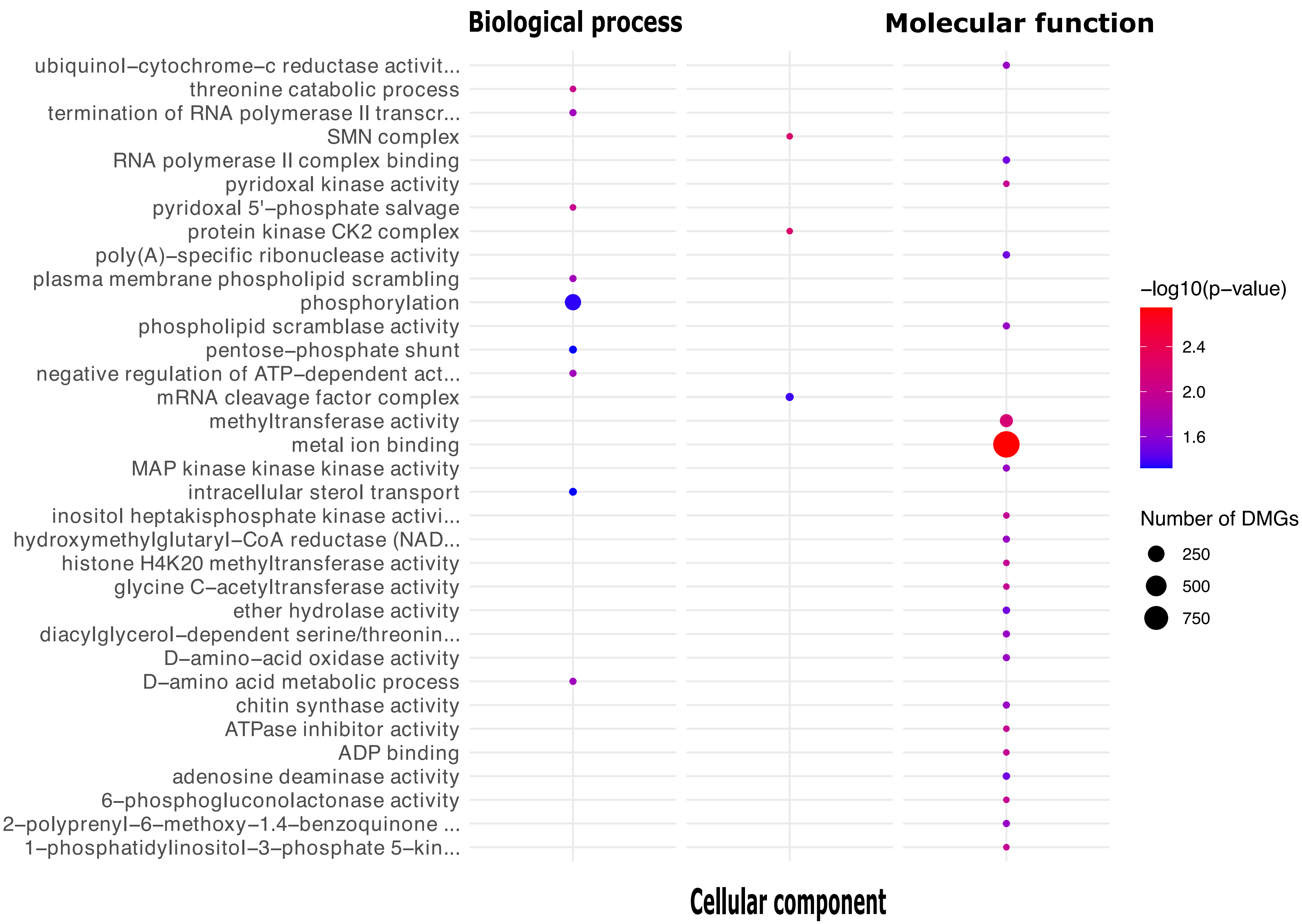

### Fig S6. DMGGotermalltreatments.pdf

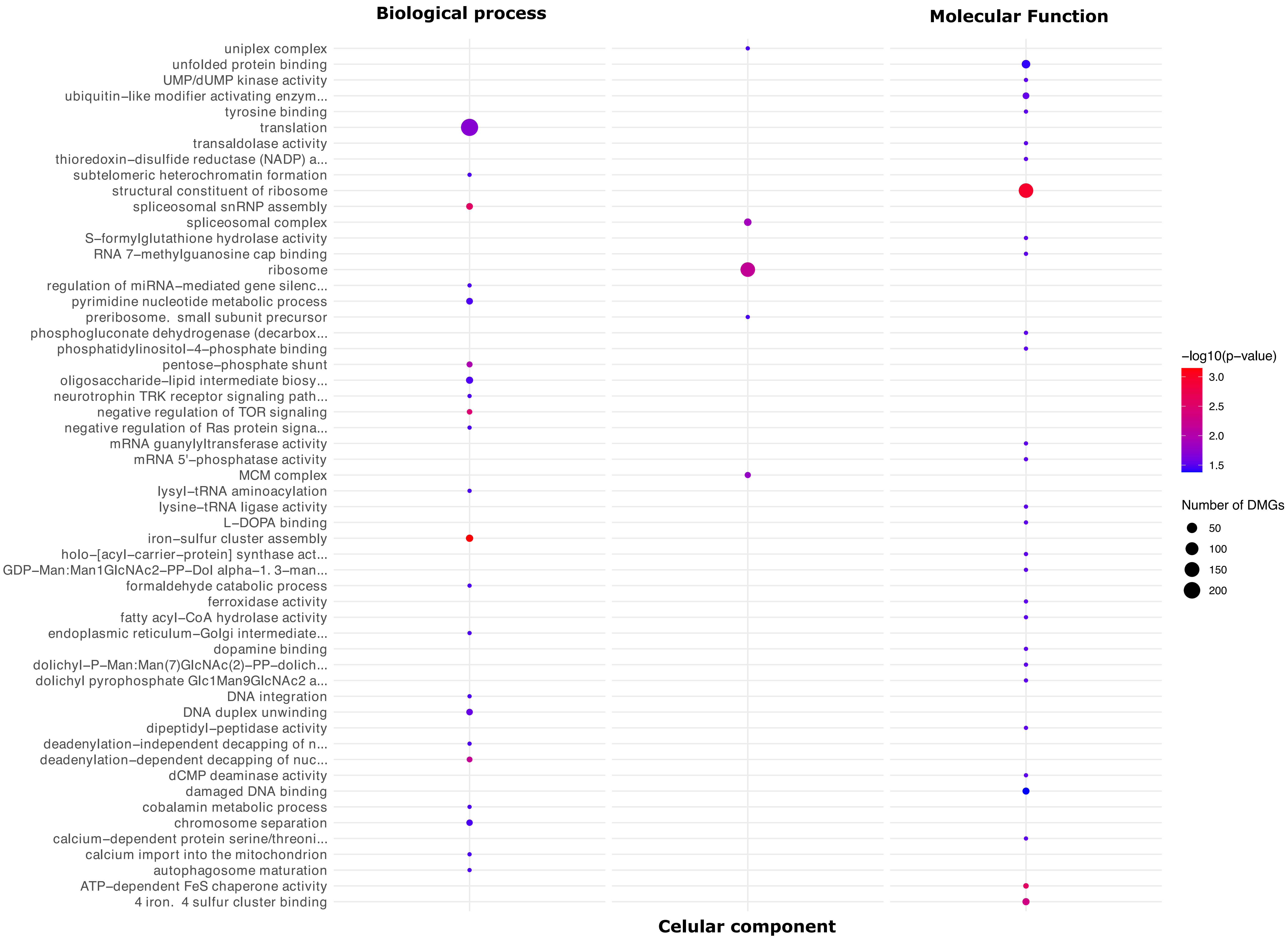

### Fig S7. DNMT methylation frequencies.pdf

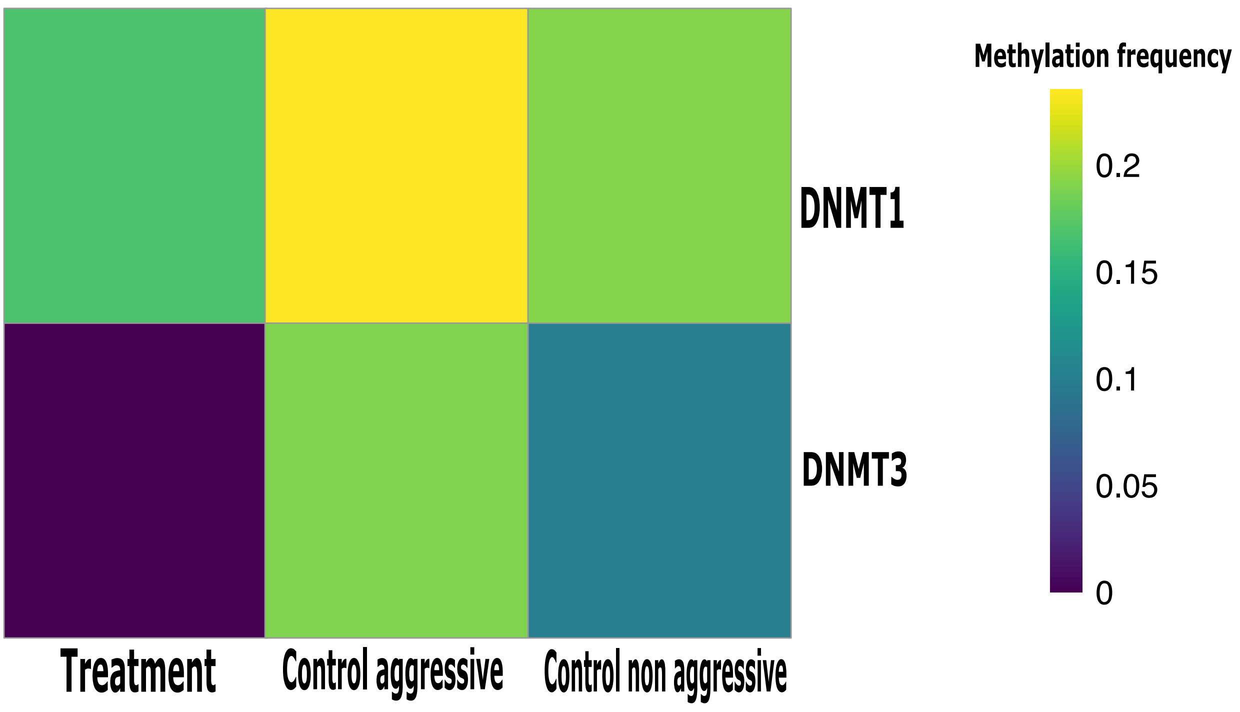

### Fig S8. Volcano CAvsCNA.pdf

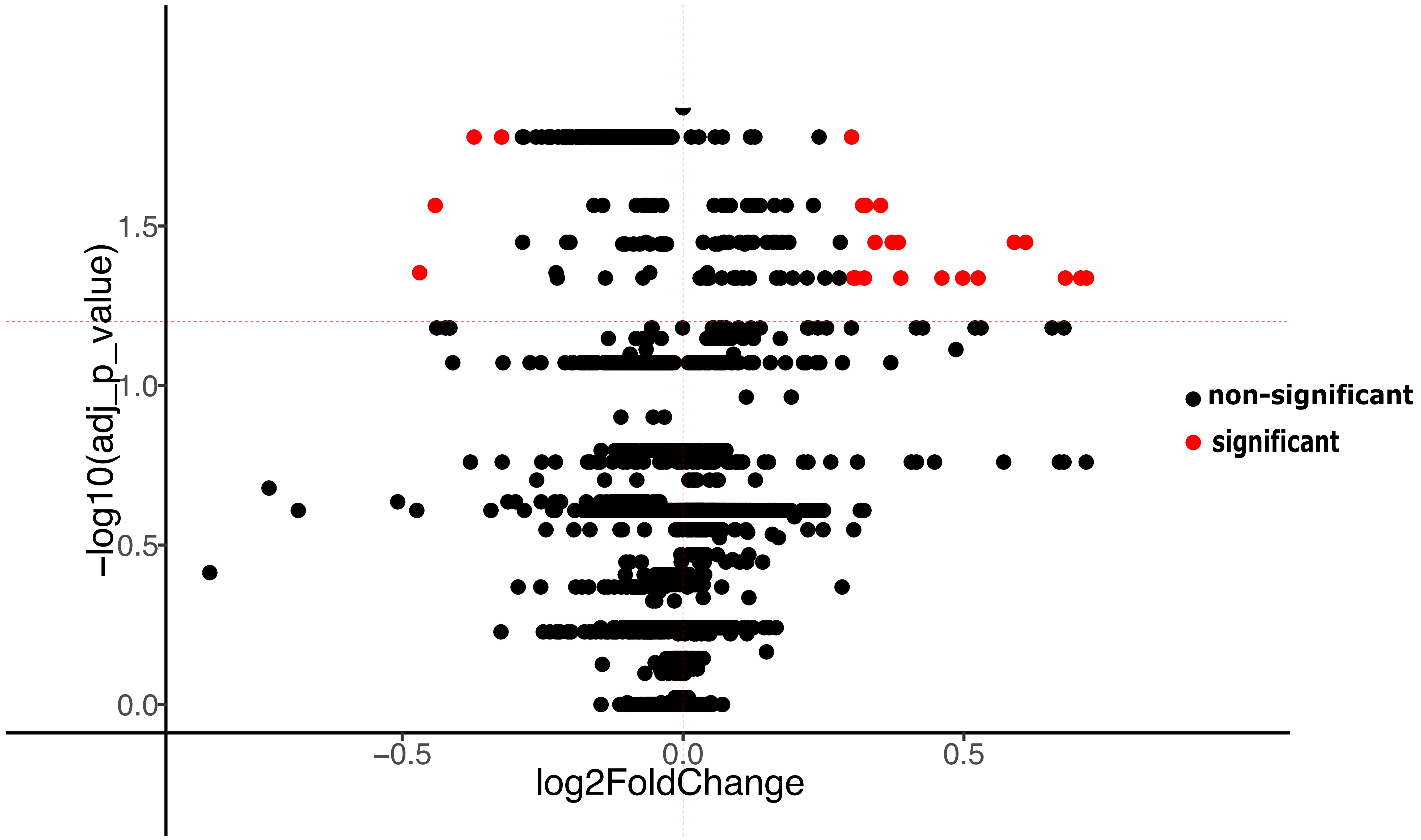

### Fig S9. Volcano control vs treatment.pdf

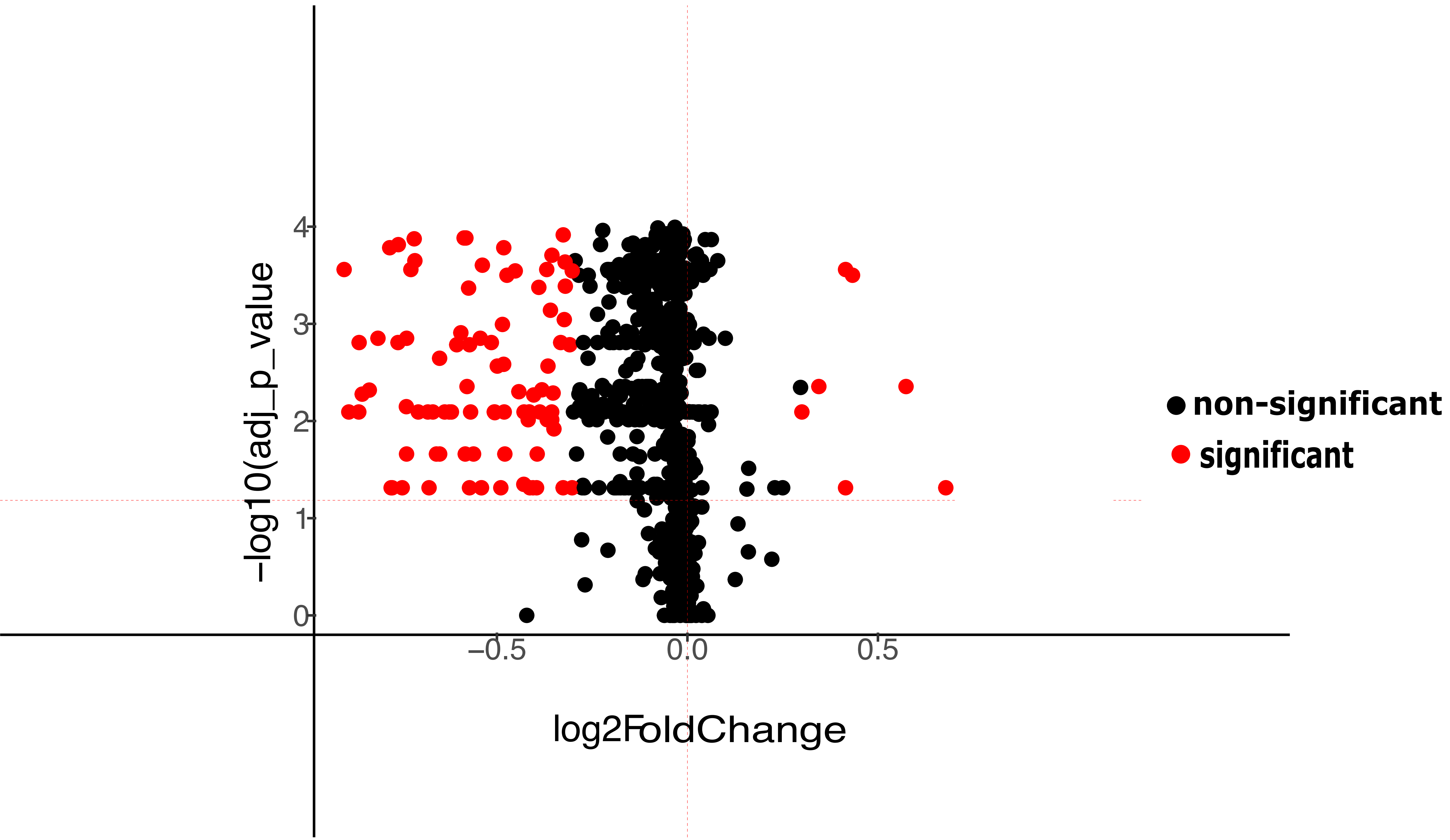

### Fig S10. CandidategenesGotermalltreat.pdf

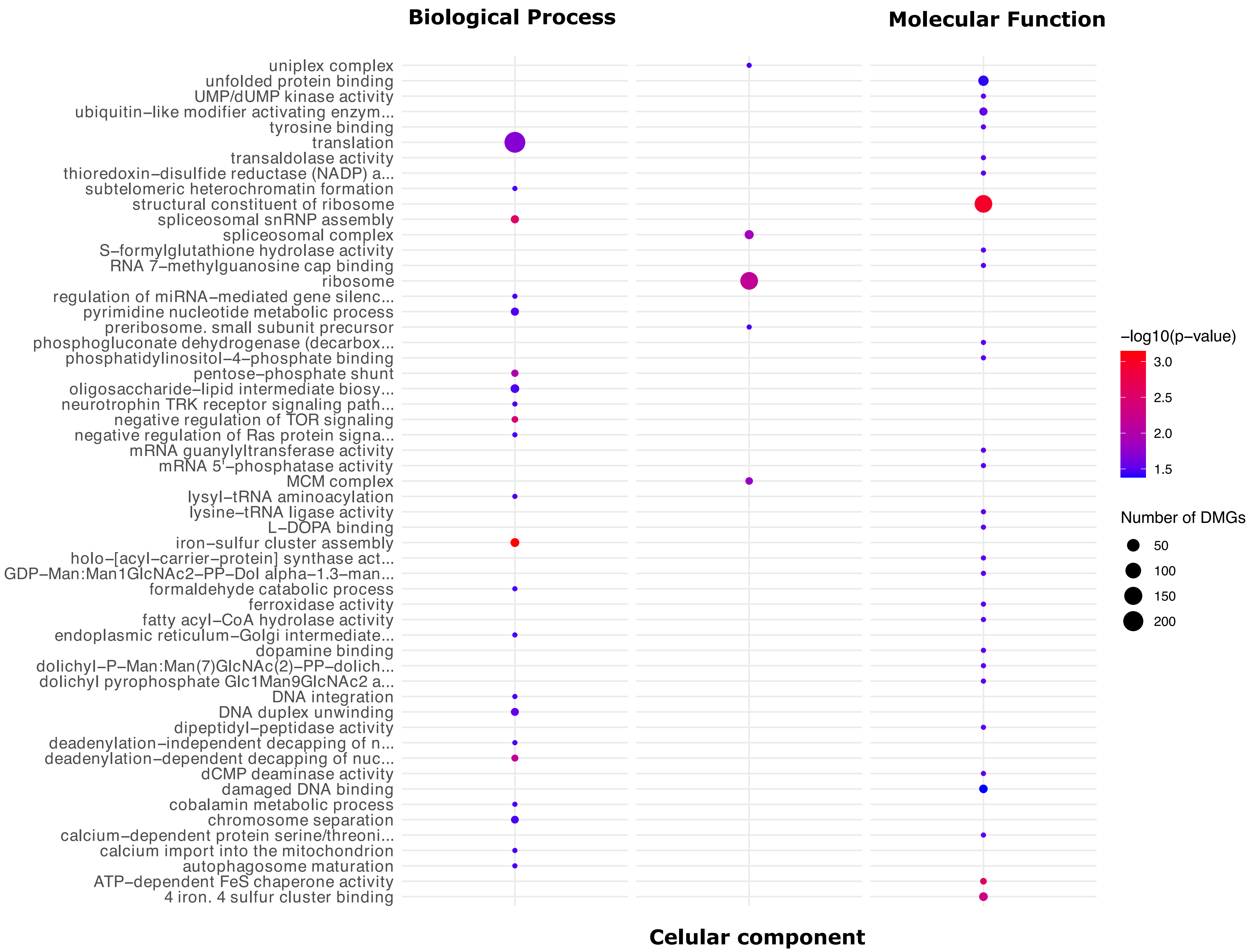

### Fig S11. Correlation20candidatesgenes.pdf

Spearman Correlation: Methylation Frequency vs. Aggression Rank

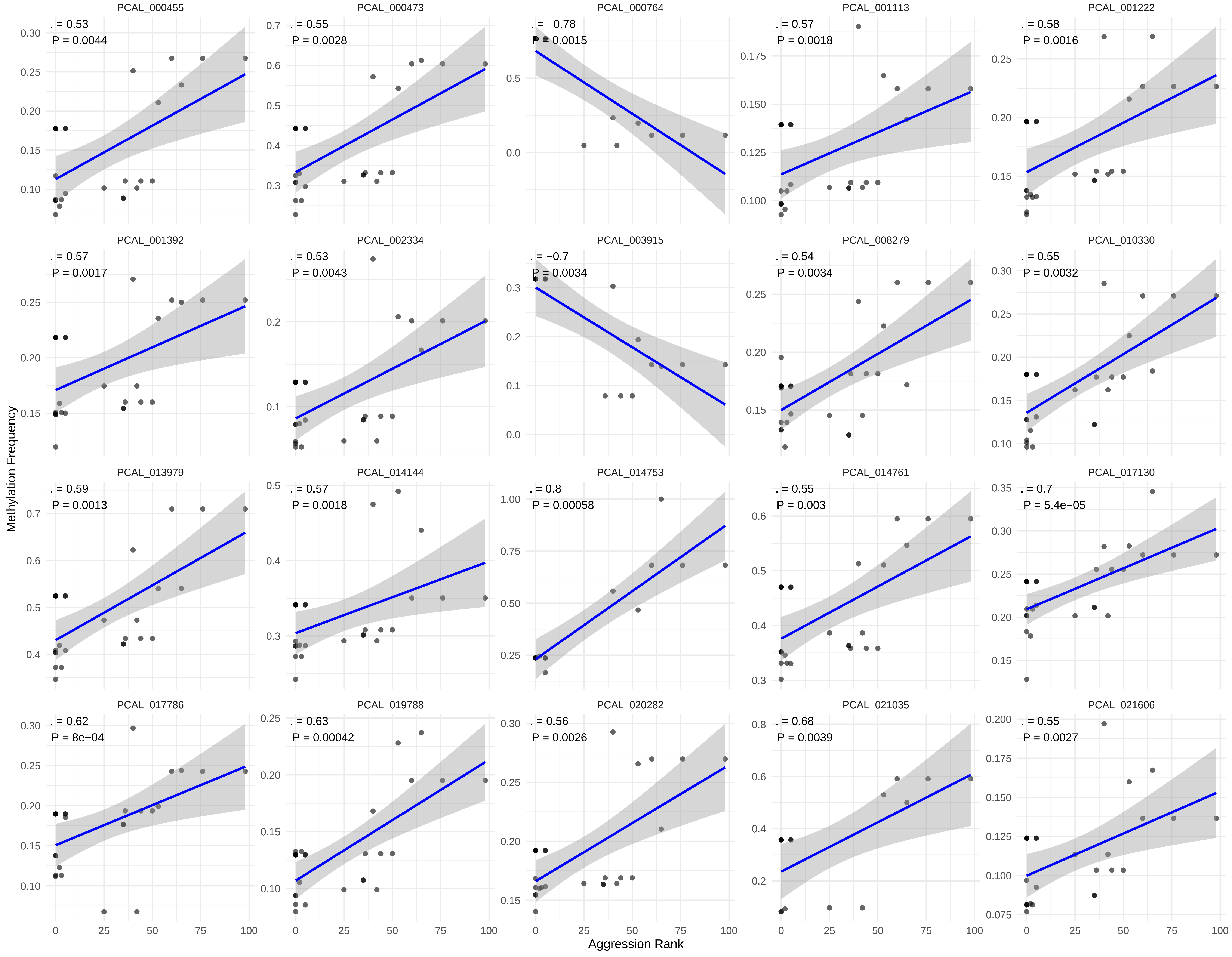

### Fig S12. weightedmethyldelayedvsnondelayed.pdf

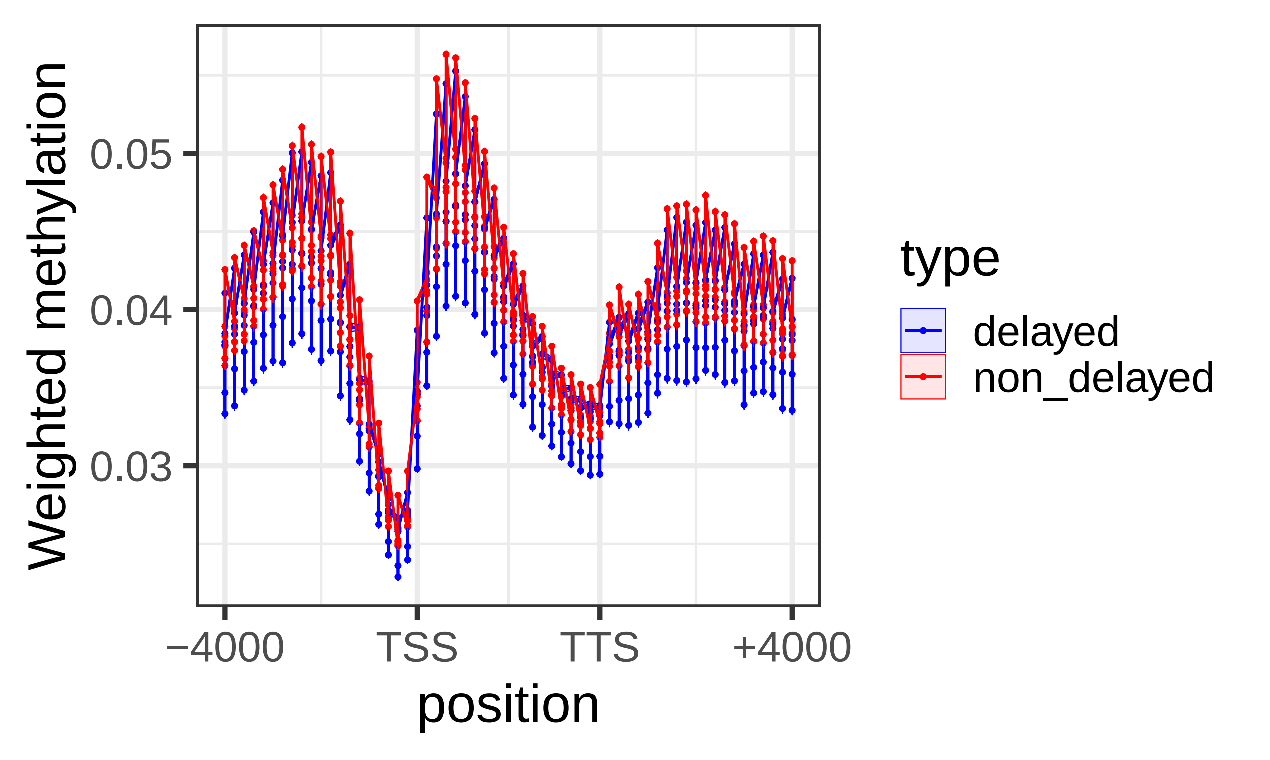

### Fig S13. VolcanoplotWGBScontrolvsdelay.pdf

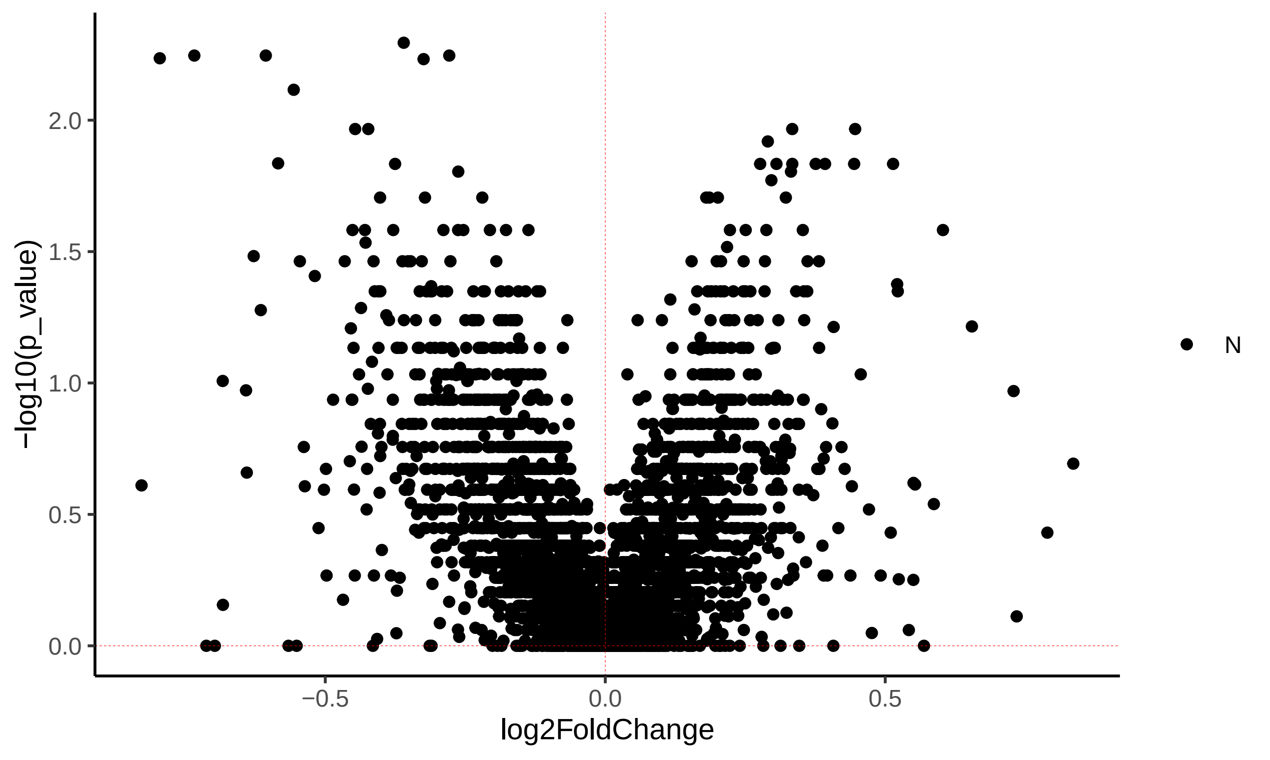
